# The retinal RNA editome is concentrated in photoreceptor-specific genes and genetically linked to vision loss

**DOI:** 10.1101/2023.06.22.543349

**Authors:** B RE Ansell, R Bonelli, S N Thomas, A Manda, R Ratnapriya, M Pinelli, A Swaroop, D diBernardo, S Banfi, M Bahlo

## Abstract

**BACKGROUND:** Conversion of adenosine in RNA to inosine by ADAR enzymes, termed ‘RNA editing’, occurs at thousands of sites across the transcriptome, and is required for healthy development of the central nervous system. RNA editing can modify protein sequences, and dampen the innate immune response. RNA editing is tissue-specific and partly genetically determined. Modifications of RNA editing sites contribute to multiple diseases, particularly neurodevelopmental and neuropsychiatric diseases. Despite the importance of RNA editing in the brain, nothing is known about this process in the human retina. We describe the landscape of retinal editing revealing its importance in key biological processes that underpin vision.

**METHODS & RESULTS:** We analysed the transcriptomes of >500 donor retinae and identified ∼153,000 high-confidence RNA editing sites. Some 80% of editing sites occurred within protein-coding RNA, with the majority in intronic *Alu* repeats, and 3’ UTR sequence. Novel retina-specific sites were concentrated in genes related to photoreceptor function and which cause retinitis pigmentosa, most notably in PDE6A. Exonic, protein recoding sites were enriched in zinc-finger domains. AMD subjects exhibit relatively few differences in RNA editing compared to controls, consistent with limited gene expression differences. We identified ∼10,000 editing QTLs. The genetic architecture of editing in the retina resembles the brain, whereas editing and expression QTLs in the retina show modest genetic overlap. We report colocalization between edQTLs and retinal disease GWAS peaks for age-related macular degeneration, glaucoma and macular telangiectasia. These findings provide new insights into epi-transcriptomic regulation of genes critical for vision, and elaborate putative genetic disease driver mechanisms that appear to be independent of changes in gene expression.

## INTRODUCTION

Conversion of adenosine to inosine in RNA by adenosine deaminase enzymes of the ADAR family, termed ‘RNA editing’, occurs at thousands of sites across the human transcriptome, and is essential for healthy central nervous system (CNS) development. RNA editing limits the immunogenicity of endogenous double stranded RNA, such as is formed by ubiquitous *Alu* repeats which constitute 11 % of the human genome (Deininger 2011). Editing of codons produces amino acid substitutions that modulate the function of neurotransmitter receptors (Nishikura 2016). In non-coding RNA, this process is evidenced to affect the rate of transcription, splicing and translation. RNA editing has been extensively documented in bulk transcriptome data from 52 healthy adult tissues including 13 brain regions (Tan et al. 2017). More recently RNA editing has begun to be described in single neurons (Ansell et al. 2021) providing novel insights into the high penetrance of editing when viewed at single cell level, which is masked by bulk data. Editing has been shown to be perturbed in neuro/psychiatric conditions including ALS, schizophrenia, autism spectrum disorder and unipolar depression (Breen et al. 2019; Tran et al. 2019; Moore et al. 2019; Weissmann et al. 2016). Recent investigations of the genetic correlates of RNA editing found a substantial component of the common genetic risk of inflammatory conditions is mediated by RNA editing differences (Li et al. 2022b). *Alu* accumulation in retinal pigment epithelial (RPE) cells is a feature of AMD geographic atrophy. *Alus* promote NLRP3 inflammasome activation leading to RPE apoptosis, and thus are likely a key driver of AMD pathogenesis (Ambati et al. 2021). Despite the recent intensive research focus on RNA editing (Minton 2022) and the association between *Alu* accumulation and AMD, this process has yet to be investigated in the retina which remains a relatively under-studied tissue of the central nervous system.

The neural retina is an outgrowth of the developing brain that contains all major CNS cell types along with specialised neuronal cells, the photoreceptors. Age-related macular degeneration and (AMD) primary open-angle glaucoma (POAG) are neurodegenerative retinal diseases with high genetic heritability as well as known environmental risk factors. Together these diseases are projected to affect 200m of the world’s people by 2040, and constitute a growing portion of the global burden of vision loss (Rein et al. 2022; Wong et al. 2014). With increasing life expectancy in low and middle income countries, these age-related diseases comprise a growing fraction of the mounting disease burden that the world faces in the coming decades. For AMD and POAG, 69 and 127 genetic risk loci are reported from GWAS, respectively (Fritsche et al. 2016; Han et al. 2020; Gharahkhani et al. 2021). Macular telangiectasia II (‘MacTel’) and retinitis pigmentosa (RP) are less common retinal neurodegenerative diseases which also entail genetic risk factors (Bonelli et al. 2021; Nishiguchi et al. 2021). As a relatively accessible CNS tissue, the retina is the subject of gene therapy trials for several inherited retinal disorders (Nuzbrokh et al. 2021; Yu-Wai-Man et al. 2020). The first FDA-approved *in vivo* gene therapy-for Leber congenital amaurosis (*RPE65*; STN 125610), targets the retinal pigment epithelium which supports photoreceptor function. Thus new therapeutic targets in the retina may be more readily translated into the clinic than for other tissues.

In contrast to DNA therapy, ADAR-based RNA therapy may provide a transient, tuneable intervention with less off-target activity (Reautschnig et al. 2022), and has promising potential in the retina (Fry et al. 2020). The development of RNA therapies for the retina will require a comprehensive understanding of the endogenous RNA editing activity of the tissue. Indeed, as a first insight, the landscape and disease associations of RNA editing in the retina need to be ascertained. Molecular quantitative trait loci (QTL) describe the association between genetic variants and a molecular trait such as RNA expression (eQTLs) or RNA editing (edQTLs). Testing all common SNPs within a genetic window for association with a single quantitative trait yields a peak of association, similar to a GWAS risk locus. Colocalizing GWAS loci with QTLs is a crucial step in establishing the molecular mechanisms mediating genetic disease risk. Therefore in order to investigate the role of RNA editing in the healthy retina, and its association with retinal diseases, we quantified editing in the neural retinal transcriptome of more than 500 post-mortem donors, including 346 with AMD (Figure 2a) which have been previously assessed for eQTLs (Ratnapriya et al. 2019). In addition to landscape and first differential editing analysis of AMD, we provide the first catalogue of edQTLs in the human retina, demonstrate the higher than anticipated importance of editing in the retina and identify RNA editing variation as the potential genetic source of six risk loci identified in GWAS for multiple retinal diseases.

**Figure 1.**
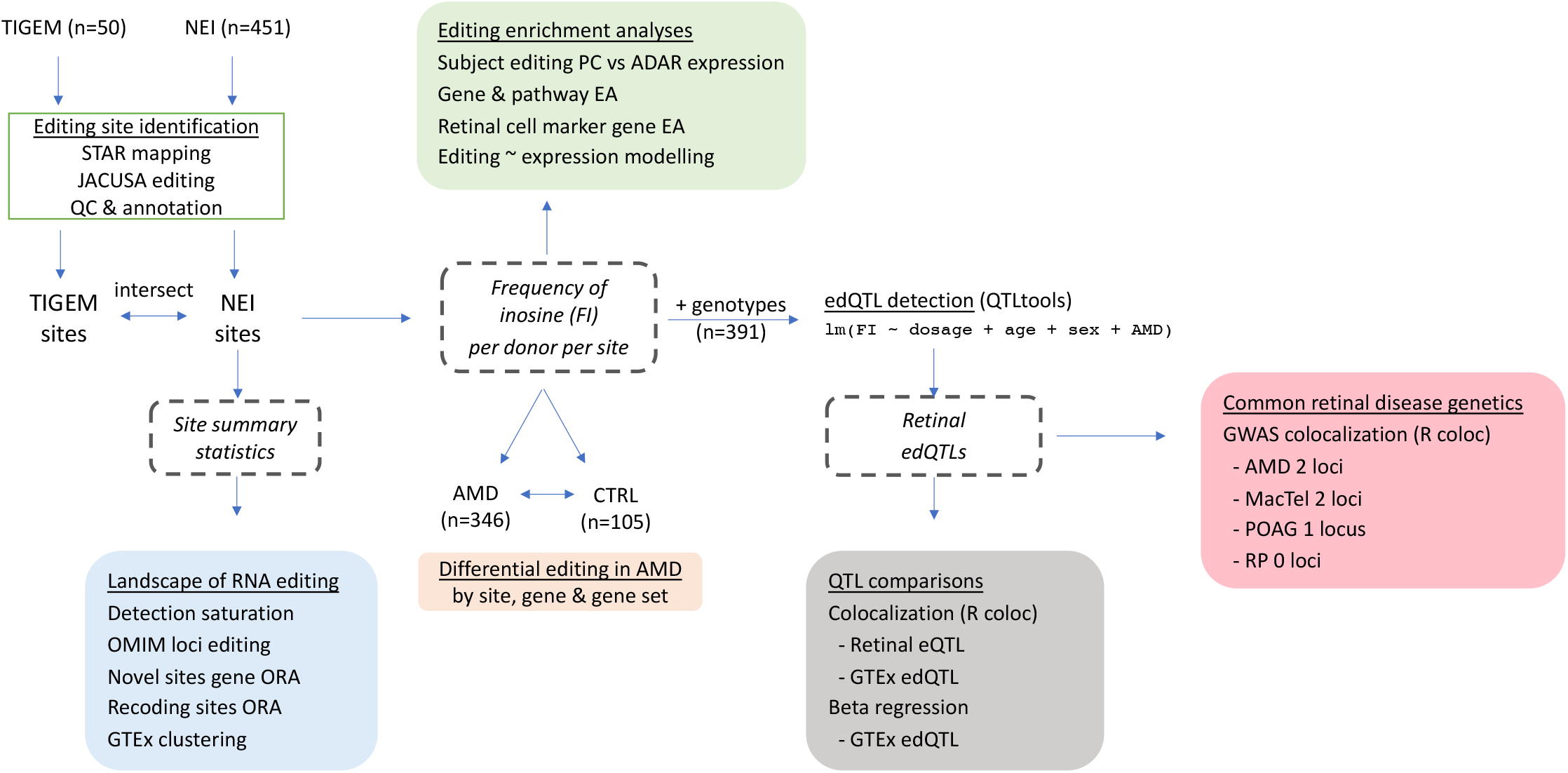
Schematic overview of retinal RNA editing analyses workflow. Preprocessing steps for Pinelli et al (TIGEM) and Ratnapriya et al (NEI) cohorts are displayed in square boxes. The key data types in dashed boxes support major analyses shown in shaded boxes. ORA: over-representation analysis; EA: enrichment analysis.

**Figure 2.**
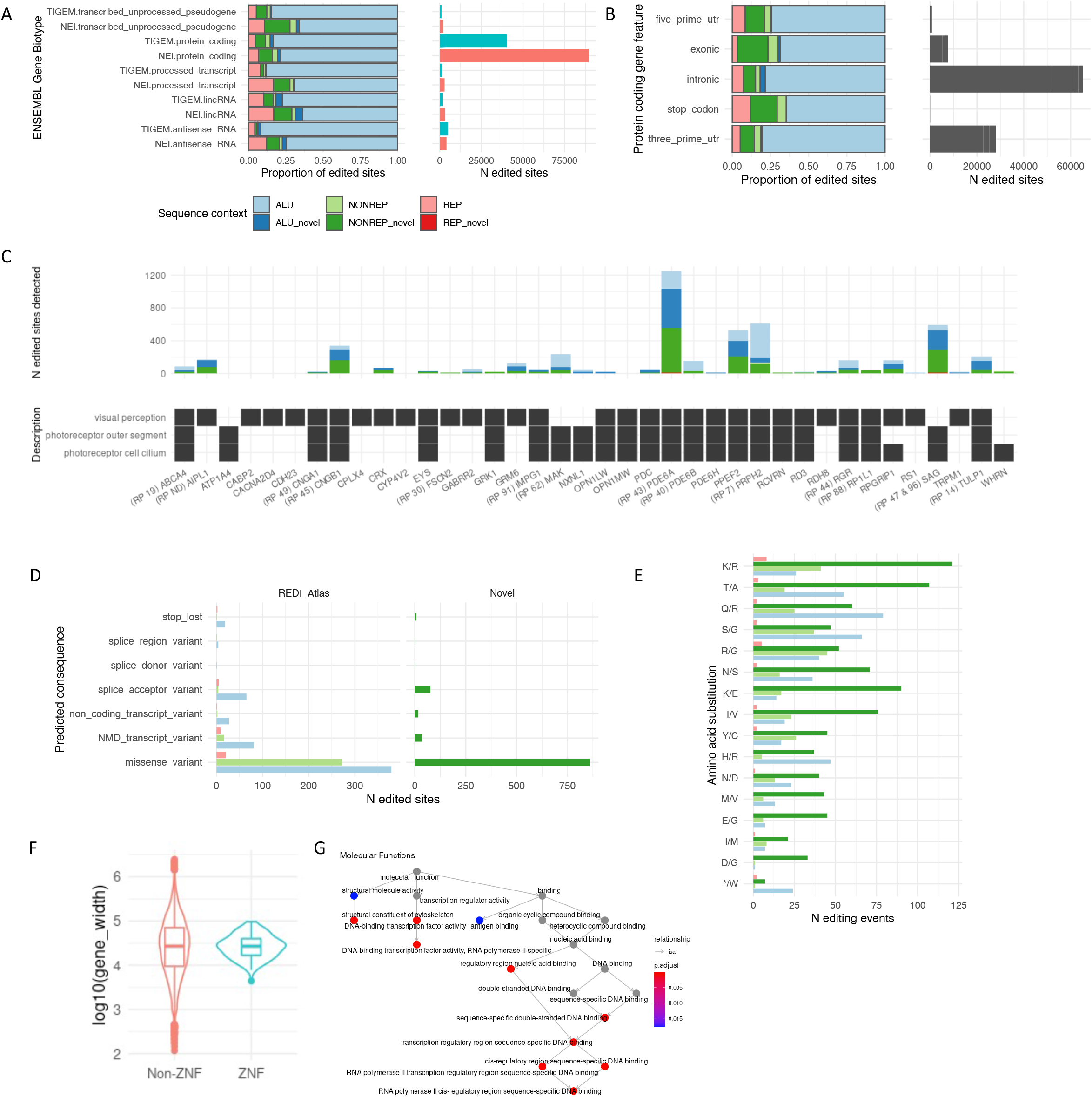
Landscape of RNA editing in the human retina. Distribution of A>I editing across the retinal transcriptome in data from independent studies by Pinelli et al (TIGEM) and Ratnapriya et al (NEI). For panels A-D, light colours denote sites previously reported in the REDIportal atlas v2.0 (Mansi et al. 2021; Picardi et al. 2017), as compared to sites likely unique to the retina and newly detected in this study (‘novel’; darker colours). A) Sequence motif context in which editing is detected is displayed as fill colours, and gene biotype is displayed on y axis. The proportion of sites in each biotype is scaled to 1 in the left panel. The absolute number of edited sites detected in each biotype is displayed at right. B) Distribution of edited sites across protein coding gene features in the larger dataset (NEI; y axis). Scaled proportions at left, with absolute number at right. C) Sum of known and novel editing sites (y axis) within three sequence contexts for genes of the GO visual perception and photoreceptor outer segment/cilium sets (grey tiles). Corresponding retinitis pigmentosa (RP) subtypes are denoted in parentheses before gene labels. D) Catalogue of predicted moderate and severe consequence of editing on gene function for sites published in the REDIportal atlas (left) and newly detected sites (right). For brevity rare consequences (predicted for <1% of dataset) are omitted. E) Catalogue of predicted amino acid (AA) substitutions induced by RNA editing. Zinc finger protein coding RNAs (ZNF) are enriched with editing-induced predicted AA substitutions. F) The relative length of ZNF genes compared to non-znf. G) Nucleic acid binding molecular functions are enriched among genes (predominantly ZNF) catalogued in D.

## RESULTS

### The landscape and predicted effects of RNA editing

RNA editing was quantified in RNAseq from 501 retinal donors using JACUSA software (Piechotta et al. 2017) (Figure 2). After quality controlling reads, editing rates were calculated as the sum of G nucleotides over all transcribed nucleotides per site for each donor (see Methods). Independent processing of cohorts published by Pinelli et al (N=50, non-AMD ‘healthy donors’) and Ratnapriya et al (N=451; 105 healthy and 346 AMD donors) yielded 46,298 unique high-confidence edited sites that were present in at least five donors in each cohort. Whereas the majority (80%) of sites detected in the smaller cohort were present in the larger cohort, an additional 106,820 sites were detected in the larger cohort. The proportion of edited sequence contexts was consistent between cohorts: most were situated within protein-coding transcripts, and 76-84% of all sites are documented within Alu repeat sequence in other human tissues (Mansi et al. 2021) (Figure 2a & S2a). Previously reported sites within non-Alu sequence constituted 7-11% of results; and novel sites in Alu sequence constituted 2-4% of the editome. We identified 13,736 novel sites lying outside of Alu repeats (9% of all sites) in the larger dataset, of which 2,283 (17%) were independently detected in the smaller cohort (Table S1). In saturation analysis, each additional donor sample added approximately 277 edited sites to the data set, and editing detection was not saturated at N=450 (Figure S1a). Having established biological reproducibility with ∼30% of the Ratnapriya et al editing sites also being detected in an independent cohort only 10% of its size, we proceeded with an in-depth analysis of the Ratnapriya et al data.

Assessment of protein-coding transcripts revealed that the majority of sites were situated within introns, followed by three prime UTRs and exons (Figure 2b). The abundance of newly detected editing sites in the retina appears to be due to extensive editing of ocular- and retina-specific genes, with 25 photoreceptor outer segment/cilium components heavily edited (Figure 2c & S2a). Accordingly, when we intersected editing sites and genomic regions associated with rare diseases in the OMIM database, transcripts arising from regions linked to retinitis pigmentosa (RP) types 7, 43, 45 and 47 were enriched with edited sites (Table S1; Figure S2b). Furthermore, the number of edited sites across all 62 loci linked to OMIM RP subtypes, was vastly higher than background (p < 10^-50^; Fig S2c). Most striking among RP-related genes was phosphodiesterase 6A (PDE6A; MIM 180071), in which 1,245 edited sites were detected, and 1,021 are novel (Figure S2d).

We detected 1,542 exonic editing sites that are predicted to modify the amino acid sequence of 668 proteins (Figure 2d & e). K>R, T>A and Q>R substitutions were the most common, and the canonical *GRIA2* Q>R site showed a FI >95% in 95% of retinal donors (Figure S1c). Approximately 56% of these protein ‘recoding’ sites have not previously been reported, highlighting retinal specificity. When grouped by gene ontology, zinc finger domain proteins which bind nucleic acid, constituted 14% (92/688) of putatively re-coded proteins. This result amounts to a 6-fold enrichment compared to the proportion of transcribed ZNF genes in the retina (<3%), and is independent of gene length (Figure 2f & g).

To examine editing in the retina relative to other human tissues, Jaccard coefficients were calculated from RNA editing sites detected in the retina and all pairs of tissues curated in the GTEx database. Unsupervised clustering of coefficients located the retina within the CNS cluster formed by multiple brain tissues and the pituitary gland. This is similar to the clustering pattern identified from gene expression analysis (Ratnapriya et al. 2019) (Figure S3a).

For quantitative analysis of site editing frequency, the proportion of edited counts (G) to all counts (A+G) was calculated for each site and donor, and reported hereafter as the frequency of inosine (FI). Principal components were constructed from FI at each site in each donor, and regressed against ADAR family gene expression. From this ADARB1 was found to correlate most strongly with the first retinal RNA editing PC (Figure S3b) demonstrating strong biological driver signals. We next curated 35 transcripts that showed exceedingly high (corrected p < 0.05) (n = 27) or low (n = 8) FI in their constituent sites relative to all edited genes in healthy donor retinae (Figure 3a). Genes with higher than normal editing included *HIF3A, MPP1, ANKH and GABRR2*, which clustered within the synaptic vesicle cycling, mitotic spindle organisation and receptor antagonist activity pathways. By contrast *SELENOW, BFAR* and *IPO9* were among transcripts that were significantly hypo-edited relative to the background transcriptome (Figure 3b).

**Figure 3.**
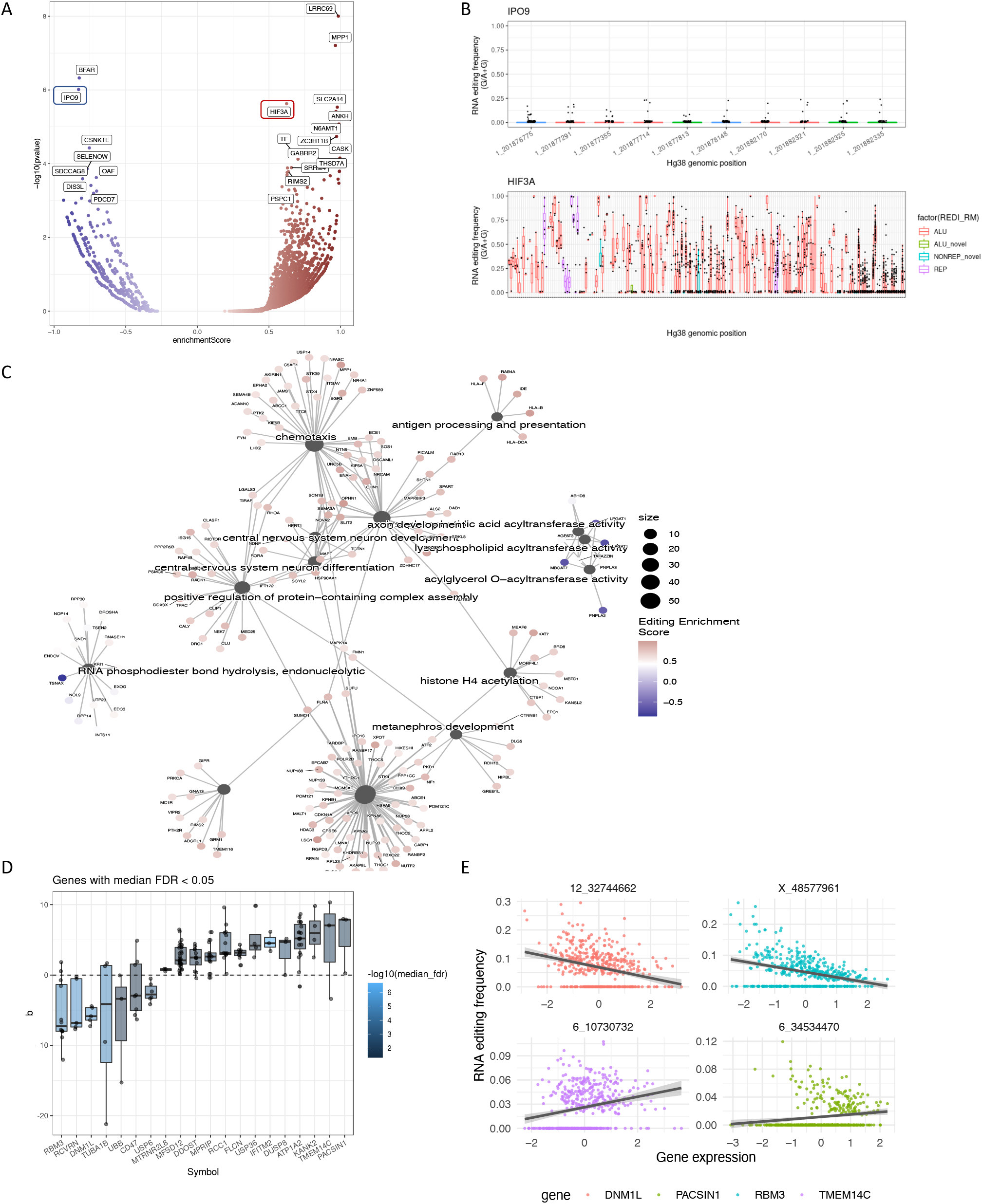
Editing enrichment analysis across the genome and editing∼expression associations. A) Volcano plot indicating gene enriched with RNA editing (reds) and hypo-edited genes (blues) relative to the average editing rate. B) Editing frequency (FI) per donor (points) at each site by genomic location (x axis) is displayed for representative hyper- and hypo-edited genes. Boxplots are coloured by sequence motif context. For simplicity HIF3A x axis labels have been omitted. C) Molecular function networks enriched among hyper-edited (reds) and hypo-edited (blues) genes. D) Genes enriched with positive and negative editing:expression associations. Each point represents the correlation (y axis) between FI for a single site and the abundance of the bounding gene (x axis). E) Representative editing:expression associations underpinning results in D, for individual sites and their bounding genes. Each point represents scaled RNA abundance (x axis) vs site editing frequency (FI; y axis) in an individual retinal donor.

When genes were ordered by mean FI and Gene Ontology (GO) enrichment testing performed, no results survived correction for multiple testing, although neuronal differentiation and post-synaptic complex genes were nominally significantly enriched with editing signal (uncorrected p < 0.05)(Figure 3c). Conversely presynaptic genes, involved in vesicle recycling, were nominally depleted of editing signal, as were RNA hydrolysis genes. A similar analysis on published marker genes for eight retinal cell-types (Menon et al. 2019) indicated relatively greater editing of bipolar cell marker genes (uncorrected p < 0.02; Figure S3).

To investigate relationships between transcript editing and abundance, we regressed the abundance of each transcript against the FI of each of its constituent editing sites. Abundance and FI were significantly associated for 2,861 sites within 791 genes. Additional filtering identified a set of 20 genes for which the median editing:expression association was significant (FDR < 0.05), indicating directionally consistent and significant associations for at least 50% of edited sites (Figure 3d & e). The gene with the most consistent negative association between editing and expression was *RBM3*, whereas *PACSIN1* which showed the strongest positive associations between site editing status and expression (Figure 3d & e).

### Disease-related differential editing

The majority of the 451 retina donors in the Ratnapriya et al cohort exhibited AMD pathology (173 early AMD ; 112 intermediate, and 61 late). Even in the presence of gross pathological changes, minimal disease-related differences in gene expression were previously reported (Ratnapriya et al. 2019). Matching these gene expression results, only 5.3% of edited sites in our data were observed in AMD donors to the exclusion of unaffected controls. This was mostly technical artefact as the relatively greater number of AMD donors increased the chance of low-prevalence sites passing our minimal threshold requiring detection in ≥ 5 donors (Methods; Figure S1d). Surprisingly however, we observed substantial differences in the amount of editing of 492 individual sites in 269 genes between AMD donors and controls (FDR<0.05) (Ansell et al. 2021; Chen et al. 2017). Only two genes were significantly enriched with AMD-related sites: *ATP1B3* was more edited in AMD; whereas *LRRC69* was hypo-edited (Figure 4a). A subsequent analysis restricted to the subset of controls with protective genotypes and cases with risk genotypes at CFH and ARMS2 respectively, produced similar results (not shown). When all genes containing at least one disease-related differently edited site were grouped by gene ontology, visual perception genes were enriched with disease-related differences, generally exhibiting greater editing in AMD retinae than controls (Figure 4b).

**Figure 4.**
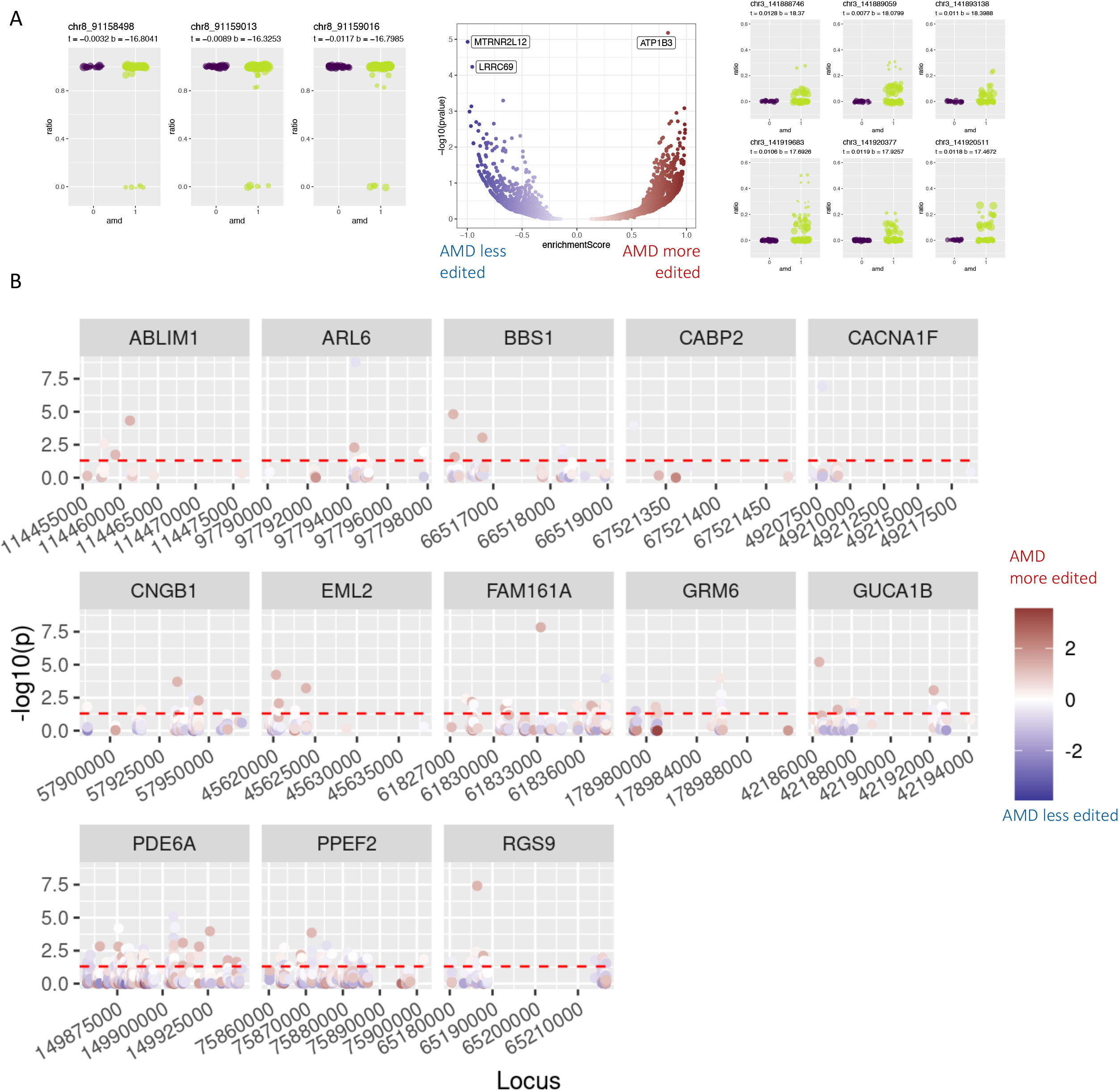
Differential RNA editing in age-related macular degeneration (AMD). Differential editing between retinae from donors with, and without age-related macular degeneration. Editing differences were determined via a counts-based GLM incorporating underlying gene expression. A) Three genes exhibited directionally consistent disease-related differences in editing. Left panel: editing frequency (FI) plots for three sites in LRRC69 that are less edited in AMD donors (x=1) than controls (x=0). Centre panel: gene-wise differential editing volcano plot. Right panel: FI plots for six sites in *ATP1B3* with increased editing in AMD compared to controls. B) Compared to background, visual function genes are enriched with differentially edited sites. Sites are represented by single points in genomic location (x axis), coloured by effect size of differential editing. Perforated line indicates the minimum threshold for significant disease association. Greater y axis height indicates greater statistical significance.

### QTL comparisons within and between tissues

Genetic variants (SNPs) that may modify RNA editing — termed ‘edQTLs’, have been curated for all tissues in the GTEx database (Park et al. 2021) and have been reported to mediate the genetic risk for several inflammatory diseases (Li et al. 2022b). We generated the edQTL dataset for the retina, based on the Ratnapriya et al cohort of 406 retinal donors for whom both genotyping and RNA was available. Fifteen donors were identified as outliers from genotype PCA analysis, and removed prior to edQTL analysis. Association tests were therefore performed between common SNPs and RNA editing (FI) using data for 391 donors, correcting for population ancestry and AMD severity as detailed in Methods. At an adjusted empirical p threshold < 0.05 we identified 9,673 edQTLs (hereafter termed ‘edSNPs’) associated with the FI of 10,475 ‘edSites’, distributed across 2,563 genes. Enrichment analysis yielded 56 protein coding genes that harboured a higher proportion of edQTLs than would be expected by chance (accounting for total number of tested sites per gene). Similarly to our AMD differential editing results, genes enriched with edQTLs were themselves enriched for visual perception functions— in particular 13 genes were identified that were highly enriched for biochemical processes in the photoreceptor cilium and outer segment (Figure 5a).

**Figure 5.**
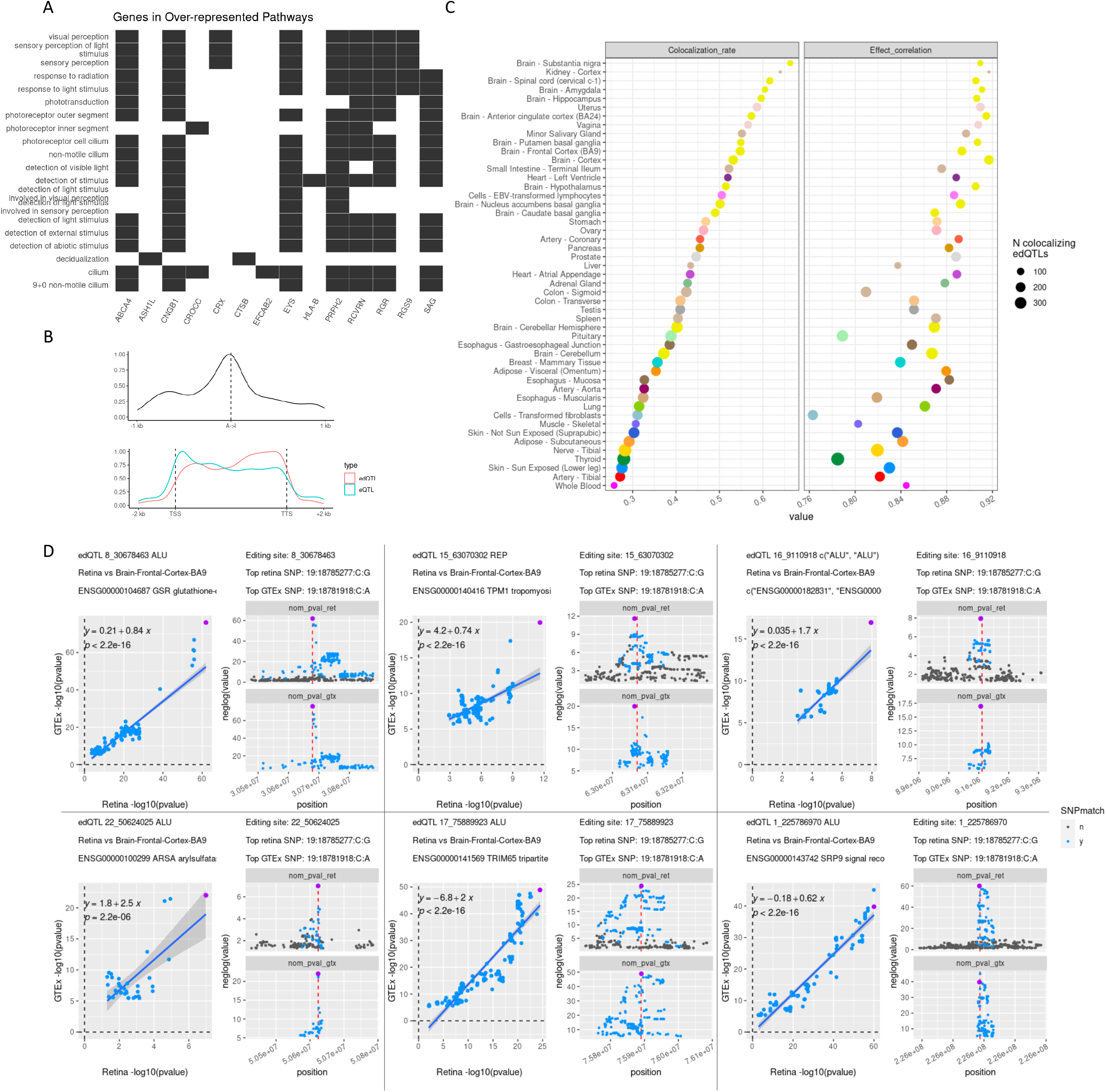
Editing quantitative trait loci (‘edQTL’) enrichment across the genome, and QTL colocalization. A) SNPs associated with RNA editing at proximal sites (edQTLs) are enriched among visual function genes, controlling for gene length. B) Editing QTLs are concentrated proximal to associated edited sites, and towards the 3’ end of protein coding genes; in contrast to eQTLs which concentrate around the transcriptional start site. C) Colocalization rate for edQTLs detected in both retina and GTEx tissues, denoted on y axis (left). Correlation in effect size for colocalized edQTLs (right), coloured by tissue of origin and sized by number of colocalized signals. D) Representative colocalization plots between retina and GTEx brain (BA9), illustrating data underlying panel C. Left panels display correlation in p values between GTex tissue (y) and retina (x); right panels display editing association peaks for retina (top) and gtex tissue (bottom).

Some 14,856 eQTLs for 10,474 eGenes are reported in the Ratnapriya et al dataset. To compare the genetic architecture of gene expression and editing in the retina, we first compared the density of eSNPs and edSNPs across protein coding gene bodies. As reported for other human tissues, eSNPs are concentrated around the transcriptional start site, whereas edSNPs show enrichment proximal to their respective edSites, and are concentrated closer to the 3’ UTR (Li et al. 2022b) (Figure 5b). We investigated 791 genes, whose editing and expression were found to be significantly correlated, for genetic associations that might drive both editing and expression signals. In total, 210 eQTL peaks (for 199 unique eGenes) overlapped with edQTL peaks for edSites that were associated with expression of the same gene. Subsequent colocalization analysis (Giambartolomei et al. 2014) identified 22 loci (11% of total) for which eQTL and edQTL genetic signals agreed (coloc PPH4 > 0.75). The majority of colocalizing e-edQTL signals pertained to editing sites and genes that were positively correlated in our preliminary analysis (Figure S3).

In contrast to the modest agreement between eQTLs and edQTLs in the same tissue, edQTL signals were strikingly similar between the retina and other human tissues generated in independent studies. Some 3,462 edSites were under putative genetic control in both the retina and at least one of the 49 other GTEx tissues. When colocalization between all pairs of overlapping edQTLS was performed, the most genetically similar tissues were cerebral brain tissues, kidney and reproductive tissues (uterus and vagina). Whole blood, tibial nerve and artery, thyroid, and sun-exposed skin were the most dissimilar to retina. Pearson correlation in the effects of colocalizing edQTLs were robust (range: 0.76-0.92) and broadly consistent with the colocalization rates (Fig 4c & d).

### edQTLs colocalize with retinal disease genetic risk

Despite the therapeutic potential of gene therapy including RNA editing for treating rare retinopathies (Maeder et al. 2019; Yu-Wai-Man et al. 2020; Fry et al. 2020), this process has not been investigated in ocular diseases to date. To investigate the extent of overlap in the genetic bases of RNA editing and ocular disease, we first curated GWAS results for AMD, POAG, RP and MacTel (Gharahkhani et al. 2021; Bonelli et al. 2021; Han et al. 2020; Nishiguchi et al. 2021). We tested for colocalization between edQTLs that fell within 0.5 Mb up- or downstream of the sentinel independent SNP(s) at each GWAS locus (Table 1). Two and four retinal edQTLs colocalized with AMD loci: chr7 *PILRA/B* (rs11771241), and chr16 *CTRB1* (rs8056814) respectively (Figure 6a). The *PILRB* edSite was intronic, however the *PILRA* edSite is predicted to introduce a missense variant causing a K106E protein substitution. Contrary to the other edQTL results in this study, the *PILRA/B* AMD GWAS locus also colocalizes with a retinal eQTL over *PILRA* (Ratnapriya et al. 2019). The four edQTL sites at the *CTRB1* locus were in fact situated within the *TMEM170A* transcript. Sites 16:75445426 and 16:75445433 were less edited with increasing dosage of the disease risk allele, whereas 16:75445570 and 16:75446123 were more edited. No retinal eQTLs are reported at this locus.

**Table 1.** edQTL and GWAS locus details for colocalizing signals.

|  | Retinal edQTL |  |  |  |  |  | edQTL | eQTL | Sequence context | Genic feature | AA substn. | GWAS |  | Coloc posterior probability of colocization |
| --- | --- | --- | --- | --- | --- | --- | --- | --- | --- | --- | --- | --- | --- | --- |
| Disease | edSite chr:bp | Edited host gene | edSNP chr:bp | Nominal p value | Slope | Adj. emp p value | N. other tissues | In retina |  |  |  | Top GWAS SNP | N. matching SNPs |  |
| AMD | 7_100374295 | PILRA | 7:100341241:G:A | 3.96E-05 | 0.37 | 5.00E-03 | 0 | y | Non-rep | exon | K106E | rs11771241 | 54 | 0.89 |
| AMD | 7_100354414 | PILRB | 7:100341241:G:A | 5.36E-05 | 0.36 | 8.00E-03 | 7 | y | ALU | intron |  |  | 35 | 0.76 |
| AMD | 16_75445426 | TMEM170A | 16:75349829:G:T | 6.43E-07 | -0.51 | 1.00E-03 | 6 | n | ALU | intron |  | rs8056814 | 29 | 0.98 |
| AMD | 16_75445433 |  | 16:75294018:A:G | 1.99E-05 | -0.42 | 2.00E-03 | 6 | n | ALU | intron |  |  | 28 | 0.93 |
| AMD | 16_75445570 |  | 16:75463548:T:C | 1.64E-06 | 0.36 | 2.00E-03 | 20 | n | ALU | intron |  |  | 59 | 0.89 |
| AMD | 16_75446123 |  | 16:75467669:A:G | 5.60E-07 | 0.42 | 1.00E-03 | 3 | n | ALU | intron |  |  | 57 | 0.86 |
| POAG | 17_47586849 | NPEPPS | 17:47703898:A:G | 2.59E-05 | 0.31 | 4.00E-03 | 0 | n | ALU | intron |  | rs1348518145 | 226 | 0.87 |
| MacTel | 17_76552668 | PRCD | 17:76699948:A:T | 1.47E-04 | 0.39 | 4.30E-02 | 0 | n | ALU | intron |  | rs11077850 | 78 | 0.66 |
| MacTel | 19_8264066 | CERS4* | 19:8263265:G:A | 3.88E-06 | -0.37 | 2.00E-03 | 0 | n | ALU | - |  | rs139412173 | 23 | 0.93 |

AMD: Age-related macular degeneration; MacTel: Macular telangiectasia II; POAG: Primary open-angle glaucoma. Full summary statistics and plots are available in supplementary materials. Number of colocizing edQTLs in other tissues filtered at $PPH4 > 0.75$ . \*4kb downstream of *CERS4*. The secondary edSNP allele is the edQTL effect allele, and increases disease risk. Adj. emp: adjusted empirical; AA substn: amino acid substitution.

**Figure 6.**
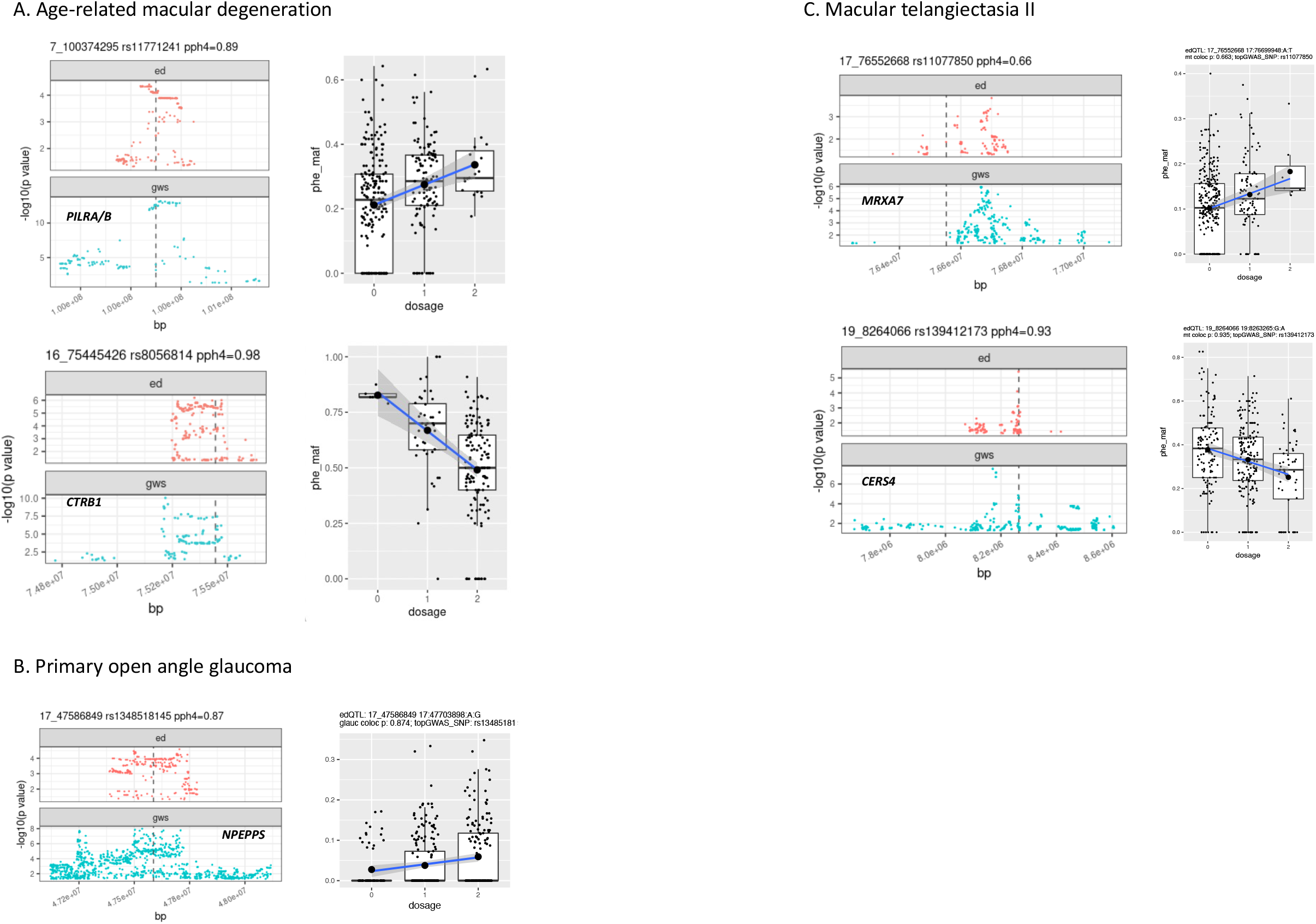
Genetic colocalization between editing QTLs and common genetic neuro-retinal disease risk. A) Colocalization in retinal edQTLs (top panel) and risk loci for age-related macular degeneration (left panels). Colocalization statistic (PPH4; 1 = perfect colocalization) is displayed above each pair. Right panels display editing frequency (FI; y axis) for individual retinal donors stratified by AMD risk allele dosage (colour) and separated by AMD disease status (x axis). B) Colocalization in retinal edQTL (top panel) and risk locus for primary open angle glaucoma (left). Right panel displays editing frequency (FI; y axis) for individual retinal donors separated by POAG risk allele dosage (x axis). C) Left: colocalization between retinal edQTLs (top panel) and GWAS risk loci for macular telangiectasia II (‘MacTel’; bottom); Right: editing frequency (FI; y axis) for individual retinal donors separated by MacTel risk allele dosage (x axis).

Published POAG GWAS peaks that colocalized with retinal edQTLs encompass haplotypes longer than 1 Mb which yield a plethora of candidate genes and genetic variants that could underpin these GWAS signals (Gharahkhani et al. 2021). Visual inspection of colocalization signals over wider genomic spans (see Methods), revealed colocalization between the POAG risk locus (rs1348518145) on chromosome 17 and an edQTL putatively affecting the FI of an intronic site in the aminopeptidase-encoding gene *NPEPPS* (Figure 6b). The single GWAS peak for RP did not colocalize with any edQTLs, however a potential protein re-coding edQTL (edSite 8:10610068; edSNP 8:10608432:T:A) was detected in the *RP1L1* gene, linked to RP subtype 88 (Bowne et al. 2003; Davidson et al. 2013). Lastly, we found two MacTel loci that colocalized with retinal edQTLs -on chromosomes 17 (rs11077850) and 19 (rs139412173) (Figure 6c). These loci proposed to impinge on the function of the matrix-remodelling associated protein 7 (*MXRA7*), and the ceramide synthesis pathway protein encoded by *CERS4* respectively (Bonelli et al. 2021; Gharahkhani et al. 2021).

## DISCUSSION

RNA editing variously affects protein sequence, RNA processing and abundance, and innate immune activation. Editing is dysregulated in multiple neuro/psychiatric illnesses, and underpins genetic risk for inflammatory diseases. However the prevalence and role of RNA editing in the retina has remained opaque. We examined the landscape of editing in 501 donor retinae, constituting the largest human CNS tissue cohort analysed to date. The approximately 150,000 sites detected in this cohort is a function of the large sample size (SFig 1d), with our resampling analysis indicating that editing detection is not complete even at this sample size (SFig 1a). Indeed, the most recent quantification of edited sites across more than 9,000 GTEx tissue samples, produced a four-fold increase compared to previous analyses of the earlier smaller cohorts (Mansi et al. 2021). Despite this massive increase, introns towards the 5’ end of transcripts are particularly under-represented in polyA-enriched RNAseq data such as is available for GTEx and the retinal cohorts reported here [Lee Law Ritchie]. As such we expect that the application of ribo-depletion protocols that preserve 5’ transcriptional regions and non-coding RNAs, would reveal many more editing sites.

Consistent with our previous work which found substantial editing-induced re-coding of RNA binding proteins in primary human neurons (Ansell et al. 2021), in the retina ZNF genes harboured an abundance of putative editing-induced amino acid substitutions. The abundance of A>I editing in transcripts encoding zinc-finger proteins is well known (Daniel et al. 2014; Shen et al. 2011). This gene family contains many human specific members, which are particularly enriched for Alu repeats, including within their exons (termed ‘Alu exonization’). The attendant concentration of ADAR activity on ZNF transcripts is demonstrated to increase the editing of proximal non-Alu sites (Daniel et al. 2014). Our finding that ZNF are enriched for editing-induced protein re-coding events within both Alu and non-Alu sequence is consistent with current understanding of the unique metabolism of this gene family, and further defines hundreds of novel re-coding events which may be unique to the retina. The extent to which these putative retina-specific events affect the activity of ZNFs as ‘master regulator’ transcription factors awaits further investigation.

Retinal genes involved in photoreceptor structure and function were particularly heavily edited in our data. Specifically, components of the cilium which connects the light-sensing outer segment to the cell body; and genes involved in membrane folding and the visual cycle (localised within the outer segment), harboured thousands of sites. The expansion of Alu repeats within opsin genes is proposed to have driven the evolution of colour vision in primates (Dulai et al. 1999). The opsin family transcripts in fact showed sparse editing compared to phosphodiesterases 6A and 6B, rod arrestin (aka *SAG*), peripherin2, the cation channel *CNGB1*, and the S/T phosphatase *PPEF2*. The latter genes include critical mediators of the rhodopsin visual cycle and are all linked to subtypes of the blinding condition retinitis pigmentosa (RP). Just as the introgression of *Alu* sequences diversified the genetics of vision-related genes, the rich layer of epitranscriptomic variation demonstrated here is likely to further augment proteomic diversity, and must now be prioritised for experimental investigation. This is especially pertinent given the vulnerability of these genes to defects that lead to blindness. Of particular interest is the extent to which the retinal organoids used to model retinopathies (Kruczek et al. 2021) recapitulate the editing landscape of the primary tissue.

Quantitative trait loci (QTLs) greatly assist in linking genetic risk loci to molecular disease mechanisms. In our data the retina closely resembles the brain in terms of both the number and location of RNA editing sites, and their genetic associations (edQTLs). This result is not surprising as the neural retina is part of the central nervous system. The overlap between retina and certain non-CNS tissues (kidney, female reproductive tissues) was unexpected at first, but likely driven by a set of conserved (non-tissue-specific) editing sites which constitute the majority of sites detected in the smaller cohorts available for those tissues. The lack of colocalization between edQTLs and eQTLs within the retina (∼15% of genomically overlapping QTLs) is similar to colocalization rates within other tissues (Park et al. 2021) and sits in stark contrast to the strong agreement between edQTLs across tissues. We attribute both technical and biological factors to this phenomenon. As mentioned above, sparse capture of 5’ introns by polyA-primed RNAseq methods hampers detection of 5’ edQTLs. Conversely eQTL search windows are centred on the (5’) transcriptional start site, and may not cover the 3’ region of long genes. This consideration aside, a recent survey by Li et al established the contribution of edQTLs to common inflammatory diseases, independent of eQTLs (Li et al. 2022b), and the two mechanisms of transcriptional regulation are likely to be truly genetically independent in the retina in many cases (discussed below).

In addition to revealing an abundance of A>I editing activity in RP genes, the Ratnapriya et al cohort allowed detailed examination of the functional genomics of common retinal diseases. The cohort included 286 donors with AMD, in which we could compare RNA editing to healthy controls. As expected, many of the hundreds of AMD-associated differentially edited sites were concentrated in genes central to visual function, where increased editing in the disease state was common. This runs counter to the trend in inflammatory diseases, where decreased editing is associated with disease risk (Li et al. 2022b). The lack of differentially edited genes in AMD is surprising given the large cohort size, although we note that AMD is a highly heterogeneous and genetically complex condition wherein consistent gene expression differences are lacking (Ratnapriya et al. 2019). Colocalization of edQTLs with AMD GWAS risk haplotypes nevertheless produced further putative mechanistic insights at the *PILRA* (chr7) and *CTRB1* loci (chr16). The signal at *PILRA/B* colocalizes with a retinal eQTL linked to increased expression of both genes (Ratnapriya et al. 2019)exm2246229; (Ratnapriya et al. 2019), and the colocalizing edQTL signal is similarly associated with greater editing of sites in both *PILRA* and *PILRB* transcripts. In myeloid cells, the former gene inhibits the immune activation cascade, whereas *PILRB* mediates inflammation (Wang et al. 2013). The robust transcription of these genes in the retina and their AMD-associated genetic architecture raises questions about their processing and function at the primary disease site. As a first step, PILRA/B protein production has yet to be directly assayed in the retina (Ahmad et al. 2018).

We detected four edQTLs colocalizing with AMD risk at the *CTRB1* locus, with two increasing editing and two decreasing editing of separate sites within transcripts encoding the endoplasmic reticulum transmembrane protein TMEM170A, which highlights the potential complexity of epitranscriptomic regulation at this locus. Of the four genes proximal to the sentinel AMD risk SNP at this locus, *CTRB1* itself is lowly expressed in bulk retinal RNA and not edited. *TMEM170A, BCAR1* and *CFDP1* are robustly expressed in retinal RNA, but none are associated with AMD via retinal eQTLs. Transcriptome-wide association analysis by Strunz and colleagues in 27 non-retinal tissues indicated that all three genes may nevertheless contribute systemically to pathogenesis via eQTL effects in other tissues (Strunz et al. 2020). Taken together, the AMD-associated changes in editing at *TMEM170A* may contribute to pathology independent of an effect on RNA abundance in the retina, and implicates TMEM170A as a potential novel retinal driver gene.

Lastly, the edQTL colocalizing at the MacTel risk locus rs11077850 (chr17) may help to further refine the driver genes at this locus. The matrix remodelling protein MXRA7 is the canonical driver gene based on intersecting retinal eQTL and TWAS results, although no retinal eQTLs survive colocalization testing at this locus (Bonelli et al. 2021). Interestingly the colocalized edQTL at this locus is associated with increased editing of a site in the photoreceptor disc component transcript *PRCD* (Progressive Rod-Cone Degeneration/RP36). In this case the edSNP lies within the MacTel risk locus bounds however *PRCD* and the corresponding edSite are beyond the locus boundaries. *PRCD* is differentially expressed in MacTel patient-derived RPE cell lines compared to unaffected controls (Eade et al. 2023). This result highlights the exciting potential for edQTL analysis to extend the inference of disease genes beyond GWAS peaks, and shed light on new potential disease mechanisms.

The results presented here indicate a predominant RNA editing signal in vision-related genes, specifically those functioning in the photoreceptor outer segment. Enrichment analysis produced weak evidence of enrichment of editing in bipolar cell marker genes, consistent with a recent study of a restricted set of non-synonymous editing sites in mouse retina (Li et al. 2022a). The relative contribution of different neuronal populations to the aggregate signal will be greatly resolved with single nucleus sequencing that samples across entire transcripts (SMRT-seq), as we recently analysed for single cortical neurons (Ansell et al. 2021). Similar investigation in primary retinal pigment epithelial cells is also required to fully resolve the landscape of epitranscriptomic modifications in retinal health and disease. This tissue supports the nutritional requirements and structural integrity of the neural retina and is under intensive investigation as a primary disease site in macular degeneration (Senabouth et al. 2022).

This study is limited in several respects. The linear modelling approach for RNA editing requires conversion of allele count data to a proportion, which can violate the model assumptions. This is a weakness of adapting eQTL software (Delaneau et al. 2017) to assess other molecular QTL types, and thus inherent in all edQTL studies published to date. A glm-based approach that works with counts data is preferable, but impractically slow for the number of tests required for a transcriptome-wide analysis. Further, molecular QTL studies are underpowered relative to GWAS, and this is especially the case for studies of rare primary retinal tissue cohorts. Future studies with larger cohorts will likely reveal further editing associations with disease, and will be particularly important for dissecting the mechanisms of rarer diseases such as RP and MacTel.

In summary, our work comprehensively characterises the RNA editome of the human neural retina, identifying thousands of putative protein re-coding events, disease-related differential editing, and putative driver mechanisms at genetic disease risk loci which broaden the potential therapeutic targets for preserving vision.

## DATA / COI / FUNDING

### The authors declare no conflicts of interest

Brendan Ansell was supported by a National Health and Medical Research Council (NHMRC) Early Career Fellowship (GNT1157776) and the Marian and EH Flack Trust via the Jack Brockhoff Foundation (GNT4495). Simon Thomas was supported by the Undergraduate Research Opportunities Program. Roberto Bonelli was supported by a Melbourne International Research Scholarship and the John and Patricia Farrant Foundation. Melanie Bahlo was supported by an Australian National Health and Medical Research Council (NHMRC) Senior Investigator Grant (GNT1195236). Brendan Ansell and Melanie Bahlo are also supported by an NHMRC Synergy Grant (GNT1181010). This work was also supported by the Victorian Government’s Operational Infrastructure Support Program and the NHMRC Independent Research Institute Infrastructure Support Scheme (IRIISS).

## METHODS

### Retinal RNA datasets

Paired-end strand-specific raw retinal RNA for 451 donors published by Ratnapriya et al was downloaded from the NCBI Gene Expression Omnibus (GSE11582). Matched genotype data for 406 donors were provided by Prof Anand Swaroop (National Eye Institute, NIH). Paired-end non-strand-specific.bam files representing retinal RNA from 50 donors published by Pinelli et al were downloaded from the EBI Array Express (E-MTAB-4377), and raw reads extracted via the bedtools bamToFastq module. All software versions are provided in Supplementary Table x. The quality of the raw reads was inspected using FastQC. To remove the adapter and low-quality sequences, reads were trimmed using Trimmomatic in paired-end mode (SLIDING WINDOW 4:15; LEADING 3; TRAILING 3; MINLEN 30). The resulting paired-end reads were aligned to the Ensembl human reference genome (Genome Build Hg38, GRCh38.97) using the STAR software in basic 2-pass mode with default parameters.

### Quantifying and filtering RNA abundance and RNA editing

Gene counts were estimated for each reference gene product using featureCounts. Gene expression was filtered via limma (Ritchie et al. 2015) using the filberByExpr function, and TMM normalisation was applied before variance stabilisation transformation via voom (Law et al. 2014) to obtain normally distributed values.

To identify putative editing sites, mapped reads were deduplicated using Picardtools mark duplicates (http://broadinstitute.github.io/picard) and then submitted to JACUSA software (Piechotta et al. 2017). The JACUSA D, S and Y arguments were employed to discard variants identified within 6 bp of the start or end of RNA reads; within 6 bp of an indel or splice site; or within 7 bp of homopolymeric read sequence.

The JacusaHelper R package was used to import variant calls with test statistic values ≥1.56 and total read coverage ≥ 10. The minimal alternate (i.e. edited) allele depth was set to 3. After confirming enrichment of A>G and T>C variants in putative RNA editing sites relative to common genomic SNPs (dbSNP build 155), the later were discarded and those A>G and T>C SNPs detected in at least 5 individual donor samples were retained for further analysis.

As the Pinelli et al dataset (E-MTAB-4377) is unstranded, categorically determining the originally edited strand is not possible. Therefore to enrich the dataset for true editing sites, after removing common genomic SNPs, we imposed further filtering as follows. Sites were only retained if they were: a) present in the REDIportal RNA editing atlas v2.0 (Mansi et al. 2021; Picardi et al. 2017), and/or b) located within non-overlapping regions of an ensemble feature, and on the cognate strand (i.e. A>G sites within a feature on the 5’ strand; T>C on the 3’ strand).

The same procedure was followed for the Ratnapriya et al dataset, with the exception that the JACUSA --RF-FIRSTSTRAND argument was employed reflecting the strand-specific sequencing protocol used, and as such only A>G variants were considered as putative editing sites.

The bedtools intersect module was used to locate previously unreported sites within genomic repeats (RepeatMasker); and to identify the genic feature (e.g. start codon, exon, intron, 3’ UTR etc) occupied by sites within protein-coding genes (Quinlan and Hall 2010). Sites with insufficient coverage were distinguished from transcribed, un-edited sites using the samtools depth module with mapping and read quality thresholds set to 20 (Li et al. 2009). Only unedited sites covered by at least 10 high-quality unique reads were retained.

For saturation analysis we took ten random draws of retinal donors of increasing cohort size (N = 25 to 450), and enumerated sites detected in at least 5 donors. The mean and standard deviation of the number of sites detected in each draw sample was displayed graphically, separated by sequence context. A linear model was fit to estimate the number of additional sites detected with each additional subject. The Variant Effect Predictor (VEP v.88) was used to predict the consequences of RNA editing-induced variation on the transcriptome and proteome. Only events with moderate or high predicted impact were taken forward for analysis.

### RNA editing landscape analysis

Statistical tests, data analysis and visualisation were performed using R v4.2.1 software and tidyverse packages (Wickham et al. 2019). Unless otherwise specified all enrichment plotting and testing were performed using the ClusterProfiler R-Package (Yu et al. 2012). Principal component analysis was performed on editing sites with average editing between 10% and 90%. Correlation between ADAR genes expression in retina and the first 5 PCs was assessed. For the set of quality-controlled editing sites, minor allele frequencies (G/A+G) representing the frequency of inosine (FI) were calculated for each site in each donor retina library.

Enrichment for newly detected editing sites not reported in the REDI-portal was performed using a two-step approach. First, genes significantly enriched with newly detected sites were detected using over-representation analysis (ORA). Only genes with at least 3 mapped editing sites were used. Second, genes significantly enriched (FDR<0.05) were used as input into a Gene Ontology ORA to detect biological mechanisms significantly overrepresented by the newly detected sites.

Editing enrichment by gene and pathway was similarly performed in two steps. A directional editing enrichment analysis was performed for each gene by using GSEA in control subjects only. The editing proportion of all sites within each gene was robustly compared with the distribution of editing proportion of other sites. Genes significantly enriched with low or high editing proportion were identified by setting a threshold of 0.05 on the FDR-adjusted p-value. Second, GSEA enrichment score for each gene was then used as input in a Gene Ontology GSEA analysis to detect biological pathways highly or lowly enriched with RNA editing.

### Cell-type specific enrichment

Association between editing at each site and gene expression was performed via linear regression. For each gene, retinal expression (CPM) was used at the dependent variable and regressed against the editing proportion of any sites mapped on it. The model was corrected for AMD status, sex, age, postmortem interval (hrs), rin, library preparation technician batch. Association significance was assessed using the ANOVA Fisher test comparing the deviance of models with and without editing proportion. P-values were then corrected for FDR using an ad-hoc Benjamini-Hochberg procedure. Genes and biological pathways enriched with co-regulation between editing and expression were identified using ORA as described above.

### Differential editing analysis

Association between editing at each and AMD status was performed by using a generalised linear regression model. A quasi-binomial family was chosen to jointly model the zero-inflated counts of edited reads and non-edited reads for each sample. Each model tested for association these counts with AMD while correcting for sex, age, postmortem interval (hours), RIN, library preparation technician and batch. Significance was assessed using the ANOVA Fisher test comparing the deviance of models with and without the AMD term. Degenerated models with outlier estimated dispersion were discarded from the results set. Genes enriched with sites whose editing affected AMD risk were identified using both ORA as well as directional GSEA. Biological pathways reflecting the role of RNA editing on AMD risk were identified by ORA using as input all genes containing at least one site significantly associated with AMD risk.

### editing-QTL analysis

#### Genotype pre-processing

Genotype data was extracted from homogenized neural retinal tissue and sequenced on the Illumina HumanCoreExome array (UM HUNT Biobank v1.0) chip as detailed in (Ratnapriya et al. 2019). Sex chromosome variants were omitted and as were SNPs deviating from Hardy-Weinberg equilibrium (P-value < 1E-6). SNPs with a Minor Allele Frequency (MAF) > 0.01 genotyping rate > 95% were retained. Imputation was performed relative to the Haplotype Reference Consortium panel (v.1.1) (McCarthy et al. 2016) via the Michigan Imputation Server (Das et al. 2016) using Eagle (version 2.4) for phasing (Loh et al. 2016), and the minimac4 method for imputation (Fuchsberger et al. 2015). Imputed data was filtered based on the following thresholds: imputation quality score > 0.3, MAF >1% and deviation from Hardy-Weinberg Equilibrium p > 1E-6, then transposed to the GRCh38 genome build via UCSC liftOver (Hinrichs et al. 2006). Principal components (PCs) were generated from donor genotypes using the QTLTools pca module (Delaneau et al. 2017). PCs were z-scaled, and donors removed if their value for any of the first five PCs for either genotype or phenotype was > 4 standard deviations from the mean.

#### QTL testing

To identify modifier SNPs (i.e., ‘edQTLs’) in *cis* with each edited site, the QTLTools *cis* module was used to regress the edited allele frequency, hereafter ‘frequency of inosine’ (FI) for all donors at each site against genotype (encoded as alternate allele dose) (Delaneau et al. 2017). The FI distribution was quantile normalized, and the following covariates were included: the first five genotype PCs, age, sex, AMD severity (Minnesota Grading System, 1: no pathology; 2 through 4: mild to severe AMD), death category, post-mortem interval and RNA integrity number. The search window was limited to 200Kb upstream and downstream of the edited site (400Kb total span). Only edQTLs with a nominal p value < 0.01 were returned. Permutation of nominal p values was performed via the QTLTools permute module to identify the sentinel SNP associated with each editing site. Only top snp:site pairs with an adjusted empirical p < 0.05 were taken forward for gene burden and colocalization analysis.

#### edQTL burden

Gene and biological pathways whose editing was significantly enriched for edQTLs were detected by ORA analysis. First, each gene was tested for enrichment of sites presenting significant edQTLs. Second, each gene significantly enriched was used as input to detect over-represented gene ontology biological pathways.

#### eQTL:edQTL comparison

To identify loci associated with both RNA abundance (eQTLs) and editing (edQTLs), we intersected the previously published retinal eQTLs with the set of edQTLs involving sites associated with changed RNA abundance. Given that sentinel SNPs for the same association signal often differ due to chance or technical factors, all SNPs with association p values < 0.01 for pairs of e/edQTL loci were intersected, and those pairs with at least 10 SNPs were retained for colocalization analysis.

#### Colocalization

Colocalization of edQTL signals with retinal eQTLs (Ratnapriya et al. 2019), edQTLs in other body tissues (Li et al. 2022b), and GWAS summary statistics, was performed using the coloc R package (Giambartolomei et al. 2014; Wallace 2020). GWAS summary statistics for diseases affecting the retina and optic nerve were downloaded from the GWAS catalogue or otherwise accessed as detailed in STable x. Refined summary statistics calculated for the AMD meta-GWAS (Han et al. 2020) were generously provided by Prof Stuart MacGregor. If necessary genomic coordinates were lifted from Hg19 to GRCh38 using liftOver (Hinrichs et al. 2006).

To identify retinal eQTLs within loci where colocalizing edQTL-GWAS signals were present, we first selected only eQTLs on genes containing edQTL sites. The distance from the top edSNP to the top eSNP was calculated. In cases where these top SNPs were within 200 kB of each other, and at least 10 matching SNPs were present between the QTL summary statistics, formal colocalization analysis was also performed.

Whereas edQTLs are calculated over predetermined short genomic windows, disease risk locus spans are highly variable. To assess edQTLs colocalizing with disease risk loci, a first pass analysis was performed using a minimum colocalization posterior probability threshold (PP.H4) of 0.65. A second round of edQTL testing was performed at candidate loci, extending the genomic window. The results were overlaid on GWAS loci and visually inspected. Results supported by a robust editing signal (FI > 0 in >= 10 donors) in at least two of three genotypes, and for which the sentinel SNPs overlapped, were prioritised.

## SFigure Legends

**Figure S1**. A) Saturation analysis for editing site detection. Number of sites of each sequence context detected in at least 5 donors (y axis) in random samples of increasing size (x axis). Points represent the mean, and error bars denote 1 standard deviation of 10 random samples. Note variance is artificially low at higher sample sizes due to increasing sample overlap. B) Editing frequency (FI) in the retina of the canonical ‘gold-standard’ *GRIA2* RNA editing site that induces a Q/R substitution in the resulting protein. Each point represents a retinal donor, separated by AMD disease status. C) Comparison of prevalence of editing sites across retinal donors with AMD (y axis) and without (x axis). Pink and red points denote cohort-specific sites, which are concentrated at the low prevalence bound. D) Correlation of sample size (x axis) and number of detected editing sites (y) axis in GTEx tissues (v6) and the retina (Ratnapriya et al 2019).

**Figure S2**. Enrichment of RNA editing sites in genes critical for photoreceptor function and in which defects cause retinitis pigmentosa (RP). A) Gene Ontology terms enriched with retina-specific (novel) RNA editing sites. Red circles indicate high statistical significance than bluer circles. B) Total number of edited sites (y axis) detected in transcripts arising from genomic regions (black points) associated with rare Mendelian diseases (OMIM database; x axis). OMIM associations for loci enriched with edited sites are displayed in boxed text. Genes within associated regions are indicated in dark red text. C) Transcripts arising from RP-associated genomic regions harbour significantly more edited sites than those from non-RP-associated regions. D) Editing frequency (FI; y axis) for 1,245 sites detected in the retina-specific gene phosphodiesterase 6A. The vast majority of detected sites are novel. Each black point represents FI for a single site from an individual donor retina.

**Figure S3**. Clustering of retina and all GTEx tissues by Jaccard coefficient (intersect of RNA editing sites divided by the union). Retina is highlighted in the dendrogram with a red line. Red box encompasses central nervous system tissues, which are on the same branch. B) First five principal components of RNA editing in each retinal donor (columns) correlated with ADAR family RNA abundance (rows).

**Figure S4**. Volcano plot indicating retinal cell type-specific gene sets enriched with RNA editing signal. BPs: bipolar cells; HC: horizontal cells; AC: amacrine cells; RGC: retinal ganglion cells.

**Figure S5**. Colocalization between RNA editing QTLs (top panels) and expression QTLs (bottom panels) for which editing:expression associations were detected. Points are coloured by positive (red) or negative (teal) correlation in QTL beta values.

## Notes

### Competing Interest Statement

The authors have declared no competing interest.

## REFERENCES

Ahmad MT, Zhang P, Dufresne C, Ferrucci L, Semba RD. 2018. The Human Eye Proteome Project: Updates on an Emerging Proteome. Proteomics 18: e1700394. http://dx.doi.org/10.1002/pmic.201700394.

Ambati M, Apicella I, Wang S-B, Narendran S, Leung H, Pereira F, Nagasaka Y, Huang P, Varshney A, Baker KL, et al. 2021. Identification of fluoxetine as a direct NLRP3 inhibitor to treat atrophic macular degeneration. Proc Natl Acad Sci U S A 118. http://dx.doi.org/10.1073/pnas.2102975118.

Ansell BRE, Thomas SN, Bonelli R, Munro JE, Freytag S, Bahlo M. 2021. A survey of RNA editing at single cell resolution links interneurons to schizophrenia and autism. RNA. http://dx.doi.org/10.1261/rna.078804.121.

Bonelli R, Jackson VE, Prasad A, Munro JE, Farashi S, Heeren TFC, Pontikos N, Scheppke L, Friedlander M, Egan CA, et al. 2021. Identification of genetic factors influencing metabolic dysregulation and retinal support for MacTel, a retinal disorder. Communications Biology 4: 1–14. https://www.nature.com/articles/s42003-021-01788-w (Accessed March 2, 2021).

Bowne SJ, Daiger SP, Malone KA, Heckenlively JR, Kennan A, Humphries P, Hughbanks-Wheaton D, Birch DG, Liu Q, Pierce EA, et al. 2003. Characterization of RP1L1, a highly polymorphic paralog of the retinitis pigmentosa 1 (RP1) gene. Mol Vis 9: 129–137. https://www.ncbi.nlm.nih.gov/pubmed/12724644.

Breen MS, CommonMind Consortium, Dobbyn A, Li Q, Roussos P, Hoffman GE, Stahl E, Chess A, Sklar P, Li JB, et al. 2019. Global landscape and genetic regulation of RNA editing in cortical samples from individuals with schizophrenia. Nature Neuroscience 22: 1402–1412. http://dx.doi.org/10.1038/s41593-019-0463-7.

Chen Y, Pal B, Visvader JE, Smyth GK. 2017. Differential methylation analysis of reduced representation bisulfite sequencing experiments using edgeR. F1000Res 6: 2055. http://dx.doi.org/10.12688/f1000research.13196.2.

Daniel C, Silberberg G, Behm M, Öhman M. 2014. Alu elements shape the primate transcriptome by cis-regulation of RNA editing. Genome Biol 15: R28. http://dx.doi.org/10.1186/gb-2014-15-2-r28.

Das S, Forer L, Schönherr S, Sidore C, Locke AE, Kwong A, Vrieze SI, Chew EY, Levy S, McGue M, et al. 2016. Next-generation genotype imputation service and methods. Nat Genet 48: 1284–1287. http://dx.doi.org/10.1038/ng.3656.

Davidson AE, Sergouniotis PI, Mackay DS, Wright GA, Waseem NH, Michaelides M, Holder GE, Robson AG, Moore AT, Plagnol V, et al. 2013. RP1L1 variants are associated with a spectrum of inherited retinal diseases including retinitis pigmentosa and occult macular dystrophy. Hum Mutat 34: 506–514. http://dx.doi.org/10.1002/humu.22264.

Deininger P. 2011. Alu elements: know the SINEs. Genome Biol 12: 236. http://dx.doi.org/10.1186/gb-2011-12-12-236.

Delaneau O, Ongen H, Brown AA, Fort A, Panousis NI, Dermitzakis ET. 2017. A complete tool set for molecular QTL discovery and analysis. Nat Commun 8: 15452. http://dx.doi.org/10.1038/ncomms15452.

Dulai KS, von Dornum M, Mollon JD, Hunt DM. 1999. The evolution of trichromatic color vision by opsin gene duplication in New World and Old World primates. Genome Res 9: 629–638. https://www.ncbi.nlm.nih.gov/pubmed/10413401.

Eade KT, Ansell BRE, Giles S, Fallon R, Harkins-Perry S, Nagasaki T, Tzaridis S, Wallace M, Mills EA, Farashi S, et al. 2023. iPSC-derived retinal pigmented epithelial cells from patients with macular telangiectasia show decreased mitochondrial function. J Clin Invest 133. http://dx.doi.org/10.1172/JCI163771.

Fritsche LG, Igl W, Bailey JNC, Grassmann F, Sengupta S, Bragg-Gresham JL, Burdon KP, Hebbring SJ, Wen C, Gorski M, et al. 2016. A large genome-wide association study of age-related macular degeneration highlights contributions of rare and common variants. Nat Genet 48: 134–143. http://dx.doi.org/10.1038/ng.3448.

Fry LE, Peddle CF, Barnard AR, McClements ME, MacLaren RE. 2020. RNA editing as a therapeutic approach for retinal gene therapy requiring long coding sequences. Int J Mol Sci 21. http://dx.doi.org/10.3390/ijms21030777.

Fuchsberger C, Abecasis GR, Hinds DA. 2015. minimac2: faster genotype imputation. Bioinformatics 31: 782–784. http://dx.doi.org/10.1093/bioinformatics/btu704.

Gharahkhani P, Jorgenson E, Hysi P, Khawaja AP, Pendergrass S, Han X, Ong JS, Hewitt AW, Segrè AV, Rouhana JM, et al. 2021. Genome-wide meta-analysis identifies 127 open-angle glaucoma loci with consistent effect across ancestries. Nat Commun 12: 1258. http://dx.doi.org/10.1038/s41467-020-20851-4.

Giambartolomei C, Vukcevic D, Schadt EE, Franke L, Hingorani AD, Wallace C, Plagnol V. 2014. Bayesian test for colocalisation between pairs of genetic association studies using summary statistics. PLoS Genet 10: e1004383. http://dx.doi.org/10.1371/journal.pgen.1004383.

Han X, Gharahkhani P, Mitchell P, Liew G, Hewitt AW, MacGregor S. 2020. Genome-wide meta-analysis identifies novel loci associated with age-related macular degeneration. J Hum Genet. http://dx.doi.org/10.1038/s10038-020-0750-x.

Hinrichs AS, Karolchik D, Baertsch R, Barber GP, Bejerano G, Clawson H, Diekhans M, Furey TS, Harte RA, Hsu F, et al. 2006. The UCSC Genome Browser Database: update 2006. Nucleic Acids Res 34: D590–8. http://dx.doi.org/10.1093/nar/gkj144.

Kruczek K, Qu Z, Gentry J, Fadl BR, Gieser L, Hiriyanna S, Batz Z, Samant M, Samanta A, Chu CJ, et al. 2021. Gene Therapy of Dominant CRX-Leber Congenital Amaurosis using Patient Stem Cell-Derived Retinal Organoids. Stem Cell Reports 16: 252–263. http://dx.doi.org/10.1016/j.stemcr.2020.12.018.

Law CW, Chen Y, Shi W, Smyth GK. 2014. voom: Precision weights unlock linear model analysis tools for RNA-seq read counts. Genome Biol 15: R29. http://dx.doi.org/10.1186/gb-2014-15-2-r29.

Li C, Shi X, Yang J, Li K, Dai L, Zhang Y, Zhou M, Su J. 2022a. Genome-wide characterization of RNA editing highlights roles of high editing events of glutamatergic synapse during mouse retinal development. Comput Struct Biotechnol J 20: 2648–2656. http://dx.doi.org/10.1016/j.csbj.2022.05.029 (Accessed June 10, 2022).

Li H, Handsaker B, Wysoker A, Fennell T, Ruan J, Homer N, Marth G, Abecasis G, Durbin R, 1000 Genome Project Data Processing Subgroup. 2009. The Sequence Alignment/Map format and SAMtools. Bioinformatics 25: 2078–2079. https://academic.oup.com/bioinformatics/article-lookup/doi/10.1093/bioinformatics/btp352.

Li Q, Gloudemans MJ, Geisinger JM, Fan B, Aguet F, Sun T, Ramaswami G, Li YI, Ma J-B, Pritchard JK, et al. 2022b. RNA editing underlies genetic risk of common inflammatory diseases. Nature. http://dx.doi.org/10.1038/s41586-022-05052-x.

Loh P-R, Palamara PF, Price AL. 2016. Fast and accurate long-range phasing in a UK Biobank cohort. Nat Genet 48: 811–816. http://dx.doi.org/10.1038/ng.3571.

Maeder ML, Stefanidakis M, Wilson CJ, Baral R, Barrera LA, Bounoutas GS, Bumcrot D, Chao H, Ciulla DM, DaSilva JA, et al. 2019. Development of a gene-editing approach to restore vision loss in Leber congenital amaurosis type 10. Nat Med 25: 229–233. http://dx.doi.org/10.1038/s41591-018-0327-9.

Mansi L, Tangaro MA, Lo Giudice C, Flati T, Kopel E, Schaffer AA, Castrignanò T, Chillemi G, Pesole G, Picardi E. 2021. REDIportal: millions of novel A-to-I RNA editing events from thousands of RNAseq experiments. Nucleic Acids Res 49: D1012–D1019. http://dx.doi.org/10.1093/nar/gkaa916.

McCarthy S, Das S, Kretzschmar W, Delaneau O, Wood AR, Teumer A, Kang HM, Fuchsberger C, Danecek P, Sharp K, et al. 2016. A reference panel of 64,976 haplotypes for genotype imputation. Nat Genet 48: 1279–1283. http://dx.doi.org/10.1038/ng.3643.

Menon M, Mohammadi S, Davila-Velderrain J, Goods BA, Cadwell TD, Xing Y, Stemmer-Rachamimov A, Shalek AK, Love JC, Kellis M, et al. 2019. Single-cell transcriptomic atlas of the human retina identifies cell types associated with age-related macular degeneration. Nat Commun 10: 4902. http://dx.doi.org/10.1038/s41467-019-12780-8.

Minton K. 2022. ZBP1 induces immunopathology caused by loss of ADAR1-mediated RNA editing. Nat Rev Immunol 22: 531. http://dx.doi.org/10.1038/s41577-022-00772-7.

Moore S, Alsop E, Lorenzini I, Starr A, Rabichow BE, Mendez E, Levy JL, Burciu C, Reiman R, Chew J, et al. 2019. ADAR2 mislocalization and widespread RNA editing aberrations in C9orf72-mediated ALS/FTD. Acta Neuropathologica 138: 49–65. http://dx.doi.org/10.1007/s00401-019-01999-w.

Nishiguchi KM, Miya F, Mori Y, Fujita K, Akiyama M, Kamatani T, Koyanagi Y, Sato K, Takigawa T, Ueno S, et al. 2021. A hypomorphic variant in EYS detected by genome-wide association study contributes toward retinitis pigmentosa. Communications Biology 4: 1–12. https://www.nature.com/articles/s42003-021-01662-9 (Accessed September 5, 2022).

Nishikura K. 2016. A-to-I editing of coding and non-coding RNAs by ADARs. Nat Rev Mol Cell Biol 17: 83–96. http://dx.doi.org/10.1038/nrm.2015.4.

Nuzbrokh Y, Ragi SD, Tsang SH. 2021. Gene therapy for inherited retinal diseases. Ann Transl Med 9: 1278. http://dx.doi.org/10.21037/atm-20-4726.

Park E, Jiang Y, Hao L, Hui J, Xing Y. 2021. Genetic variation and microRNA targeting of A-to-I RNA editing fine tune human tissue transcriptomes. Genome Biol 22: 77. http://dx.doi.org/10.1186/s13059-021-02287-1.

Picardi E, D’Erchia AM, Lo Giudice C, Pesole G. 2017. REDIportal: a comprehensive database of A-to-I RNA editing events in humans. Nucleic Acids Res 45: D750–D757. http://dx.doi.org/10.1093/nar/gkw767.

Piechotta M, Wyler E, Ohler U, Landthaler M, Dieterich C. 2017. JACUSA: site-specific identification of RNA editing events from replicate sequencing data. BMC Bioinformatics 18: 1586. http://bmcbioinformatics.biomedcentral.com/articles/10.1186/s12859-016-1432-8.

Quinlan AR, Hall IM. 2010. BEDTools: a flexible suite of utilities for comparing genomic features. Bioinformatics 26: 841–842. http://dx.doi.org/10.1093/bioinformatics/btq033.

Ratnapriya R, Sosina OA, Starostik MR, Kwicklis M, Kapphahn RJ, Fritsche LG, Walton A, Arvanitis M, Gieser L, Pietraszkiewicz A, et al. 2019. Retinal transcriptome and eQTL analyses identify genes associated with age-related macular degeneration. Nat Genet 51: 606–610. http://dx.doi.org/10.1038/s41588-019-0351-9.

Reautschnig P, Wahn N, Wettengel J, Schulz AE, Latifi N, Vogel P, Kang T-W, Pfeiffer LS, Zarges C, Naumann U, et al. 2022. CLUSTER guide RNAs enable precise and efficient RNA editing with endogenous ADAR enzymes in vivo. Nat Biotechnol. http://dx.doi.org/10.1038/s41587-021-01105-0.

Rein DB, Wittenborn JS, Burke-Conte Z, Gulia R, Robalik T, Ehrlich JR, Lundeen EA, Flaxman AD. 2022. Prevalence of Age-Related Macular Degeneration in the US in 2019. JAMA Ophthalmol 140: 1202–1208. http://dx.doi.org/10.1001/jamaophthalmol.2022.4401.

Ritchie ME, Phipson B, Wu D, Hu Y, Law CW, Shi W, Smyth GK. 2015. limma powers differential expression analyses for RNA-sequencing and microarray studies. Nucleic Acids Res 43: e47–e47. https://academic.oup.com/nar/article/43/7/e47/2414268 (Accessed March 18, 2022).

Senabouth A, Daniszewski M, Lidgerwood GE, Liang HH, Hernández D, Mirzaei M, Keenan SN, Zhang R, Han X, Neavin D, et al. 2022. Transcriptomic and proteomic retinal pigment epithelium signatures of age-related macular degeneration. Nat Commun 13: 4233. http://dx.doi.org/10.1038/s41467-022-31707-4.

Shen S, Lin L, Cai JJ, Jiang P, Kenkel EJ, Stroik MR, Sato S, Davidson BL, Xing Y. 2011. Widespread establishment and regulatory impact of Alu exons in human genes. Proceedings of the National Academy of Sciences 108: 2837–2842. https://www.pnas.org/doi/abs/10.1073/pnas.1012834108.

Strunz T, Lauwen S, Kiel C, International AMD Genomics Consortium (IAMDGC), Hollander A den, Weber BHF. 2020. A transcriptome-wide association study based on 27 tissues identifies 106 genes potentially relevant for disease pathology in age-related macular degeneration. Sci Rep 10: 1584. http://dx.doi.org/10.1038/s41598-020-58510-9.

Tan MH, Li Q, Shanmugam R, Piskol R, Kohler J, Young AN, Liu KI, Zhang R, Ramaswami G, Ariyoshi K, et al. 2017. Dynamic landscape and regulation of RNA editing in mammals. Nature 550: 249–254. http://dx.doi.org/10.1038/nature24041.

Tran SS, Jun H-I, Bahn JH, Azghadi A, Ramaswami G, Van Nostrand EL, Nguyen TB, Hsiao Y-HE, Lee C, Pratt GA, et al. 2019. Widespread RNA editing dysregulation in brains from autistic individuals. Nat Neurosci 22: 25–36. http://dx.doi.org/10.1038/s41593-018-0287-x.

Wallace C. 2020. Eliciting priors and relaxing the single causal variant assumption in colocalisation analyses. PLoS Genet 16: e1008720. http://dx.doi.org/10.1371/journal.pgen.1008720.

Wang J, Shiratori I, Uehori J, Ikawa M, Arase H. 2013. Neutrophil infiltration during inflammation is regulated by PILRα via modulation of integrin activation. Nat Immunol 14: 34–40. http://dx.doi.org/10.1038/ni.2456.

Weissmann D, van der Laan S, Underwood MD, Salvetat N, Cavarec L, Vincent L, Molina F, Mann JJ, Arango V, Pujol JF. 2016. Region-specific alterations of A-to-I RNA editing of serotonin 2c receptor in the cortex of suicides with major depression. Transl Psychiatry 6: e878. http://dx.doi.org/10.1038/tp.2016.121.

Wickham H, Averick M, Bryan J, Chang W, McGowan L, François R, Grolemund G, Hayes A, Henry L, Hester J, et al. 2019. Welcome to the tidyverse. J Open Source Softw 4: 1686. https://joss.theoj.org/papers/10.21105/joss.01686.

Wong WL, Su X, Li X, Cheung CMG, Klein R, Cheng C-Y, Wong TY. 2014. Global prevalence of age-related macular degeneration and disease burden projection for 2020 and 2040: a systematic review and meta-analysis. Lancet Glob Health 2: e106–16. http://dx.doi.org/10.1016/S2214-109X(13)70145-1.

Yu G, Wang L-G, Han Y, He Q-Y. 2012. clusterProfiler: an R package for comparing biological themes among gene clusters. OMICS: A Journal of Integrative Biology 16: 284–287. http://dx.doi.org/10.1089/omi.2011.0118.

Yu-Wai-Man P, Newman NJ, Carelli V, Moster ML, Biousse V, Sadun AA, Klopstock T, Vignal-Clermont C, Sergott RC, Rudolph G, et al. 2020. Bilateral visual improvement with unilateral gene therapy injection for Leber hereditary optic neuropathy. Sci Transl Med 12. http://dx.doi.org/10.1126/scitranslmed.aaz7423.

