## Supplementary figures and images for "The retinal RNA editome is concentrated in photoreceptor-specific genes and genetically linked to vision loss"

### Supplementary Figures

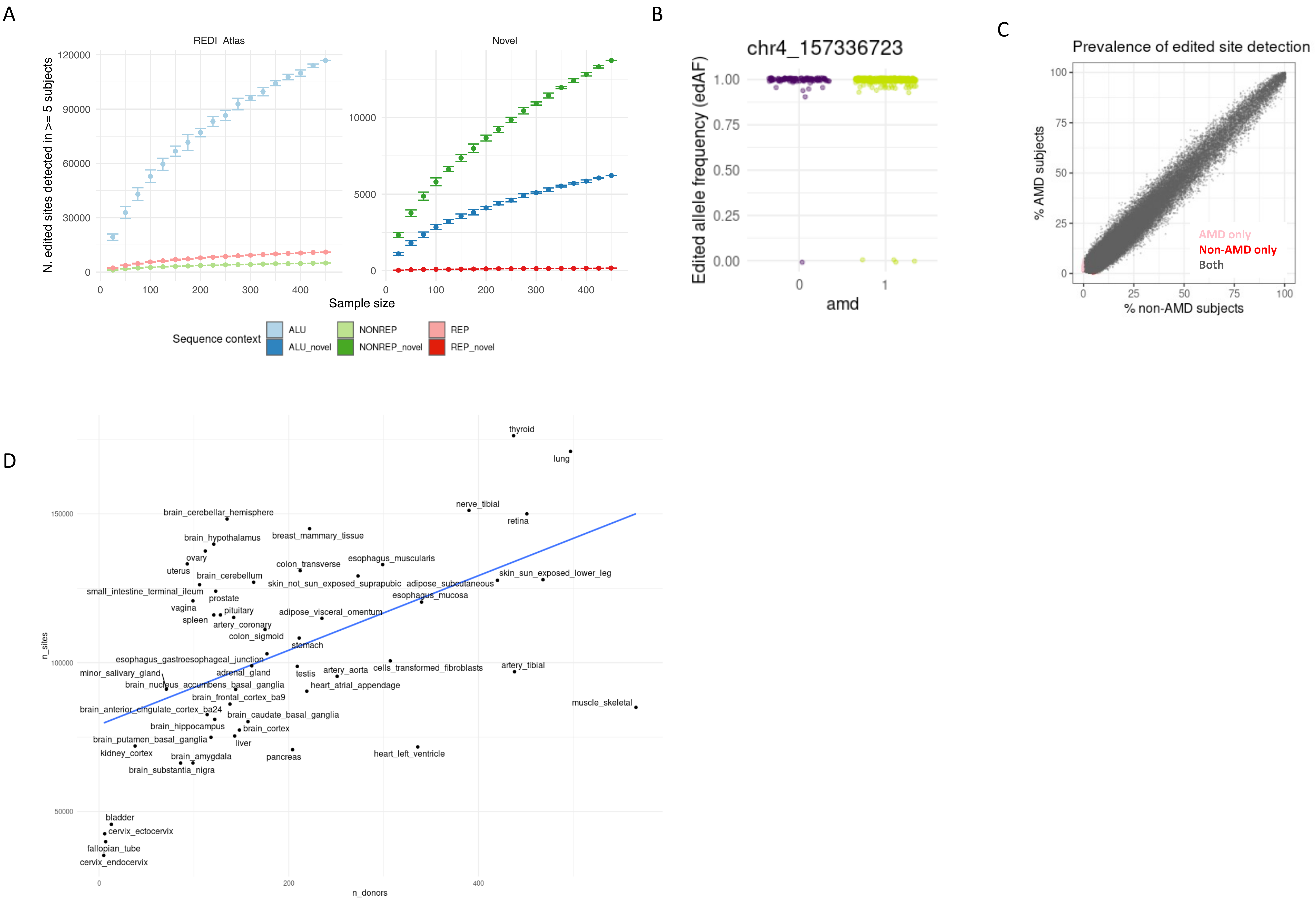

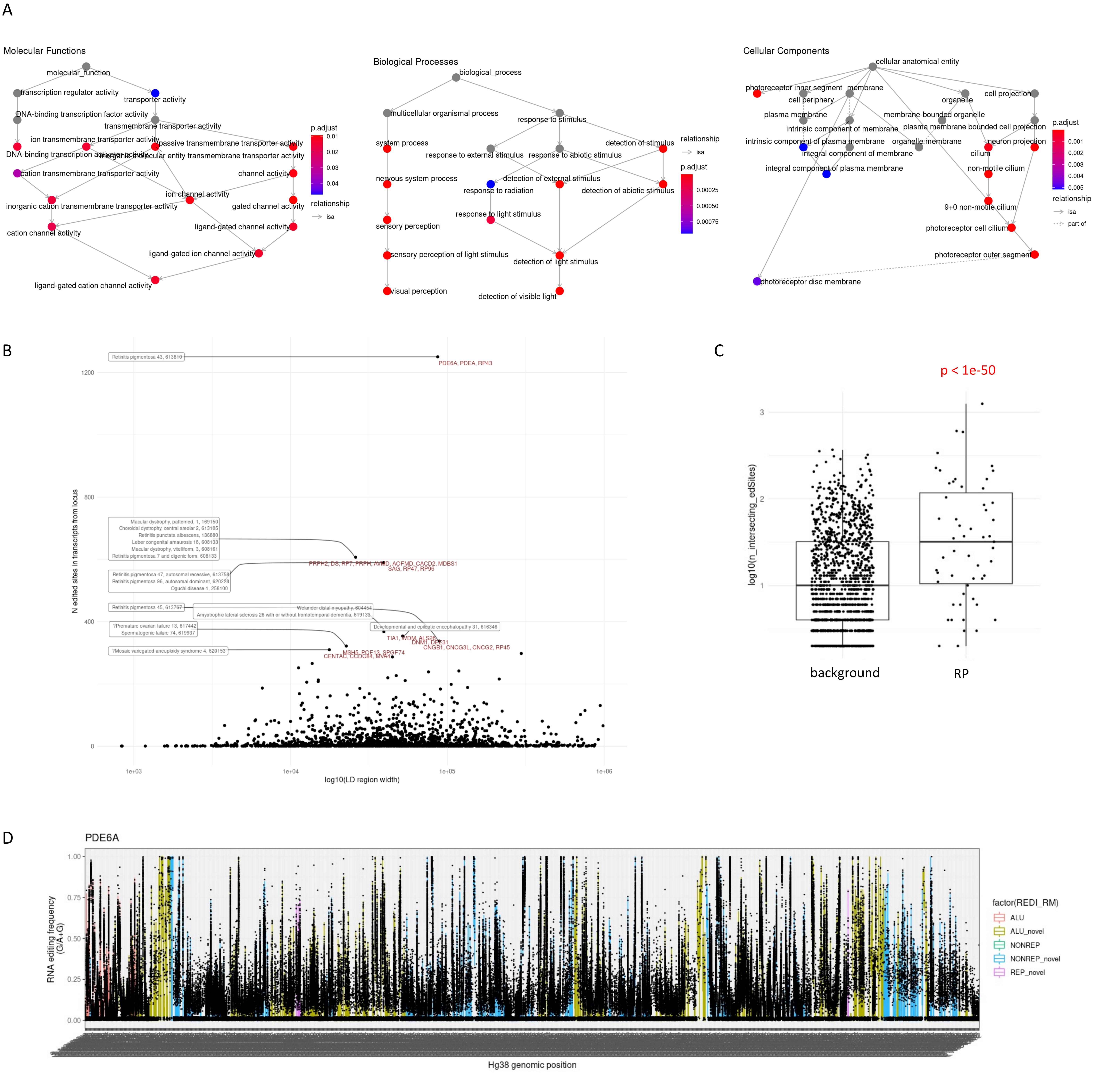

Fig S3

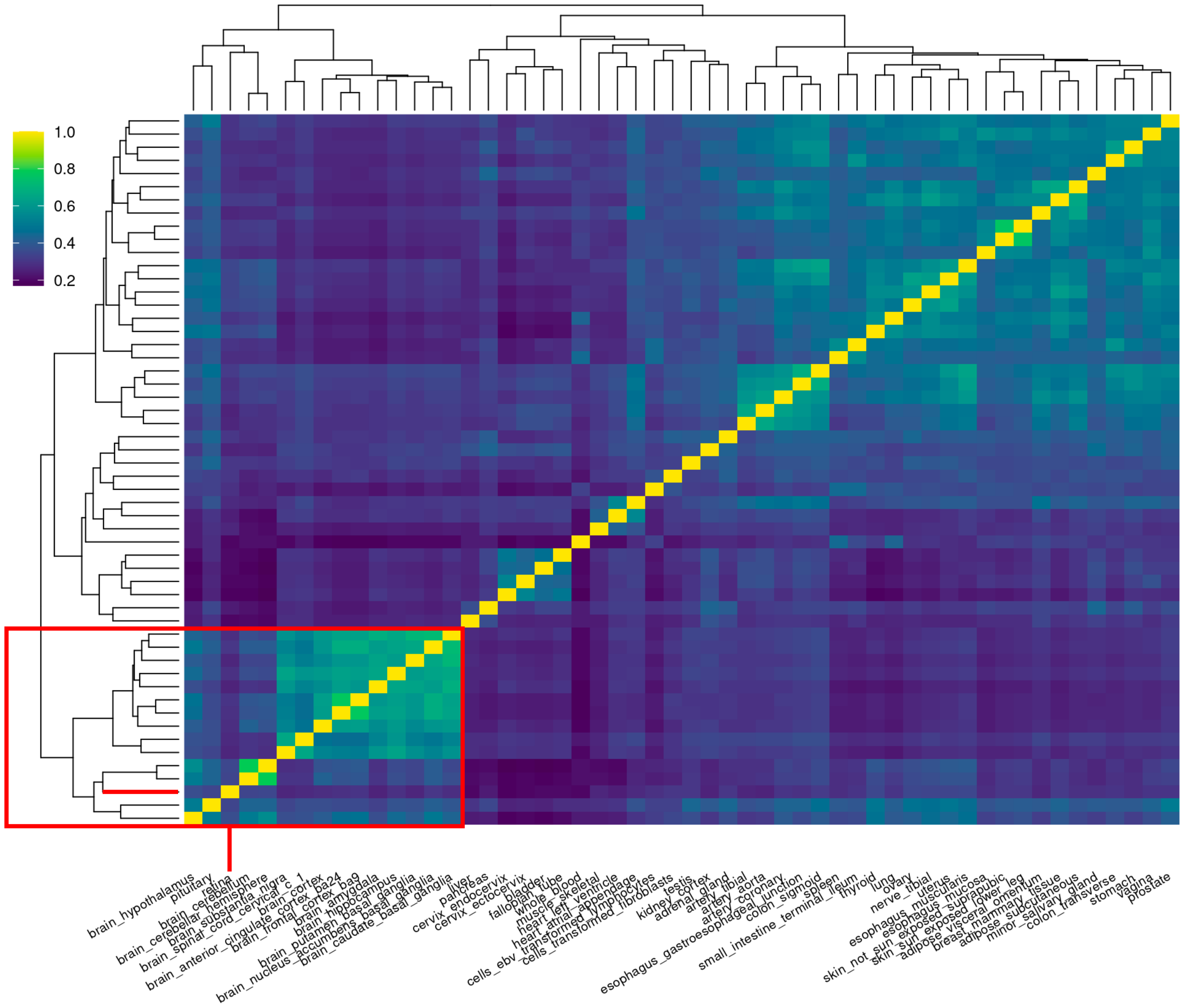


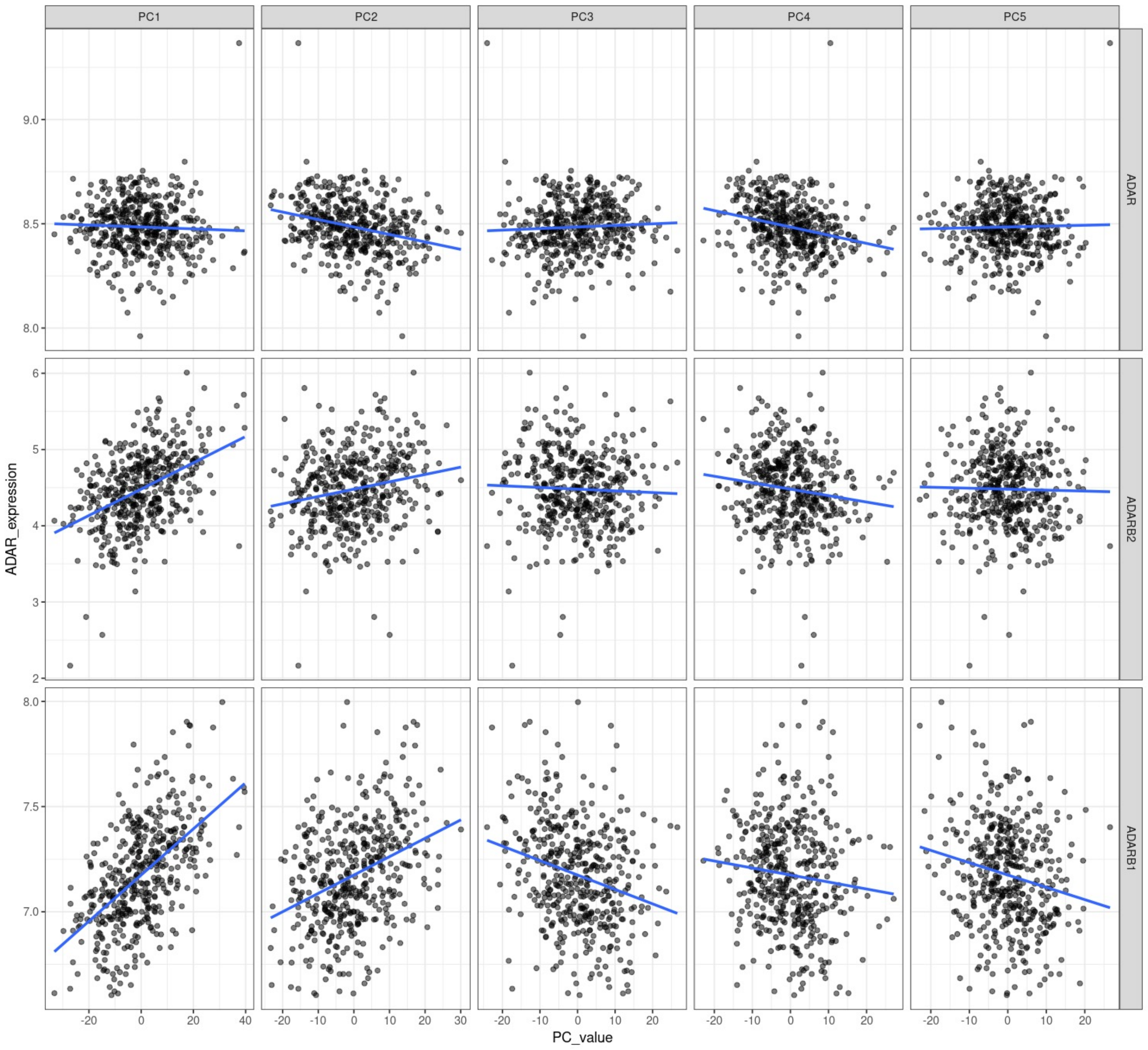


ADAR1

ADARB2

ADARB1

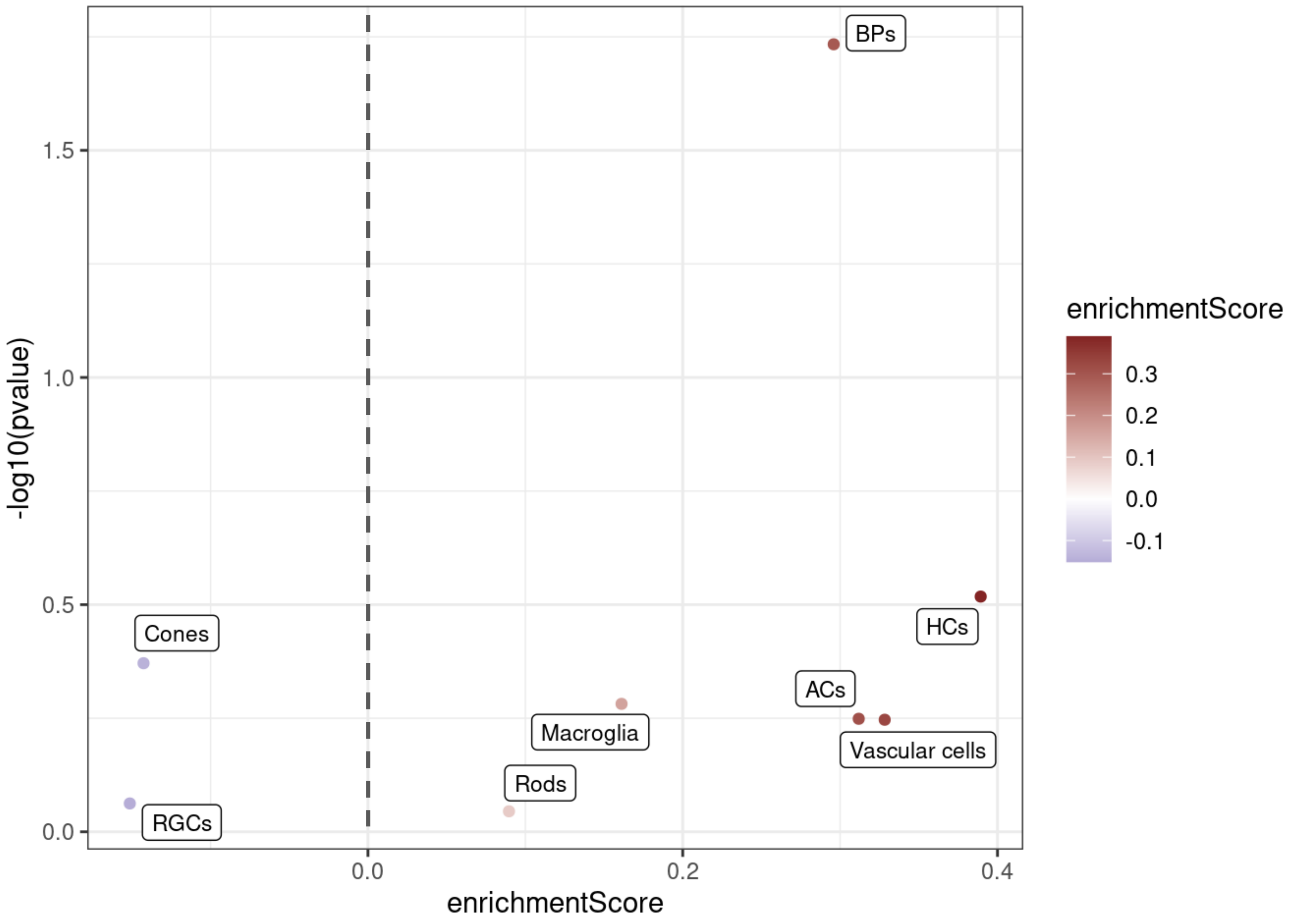

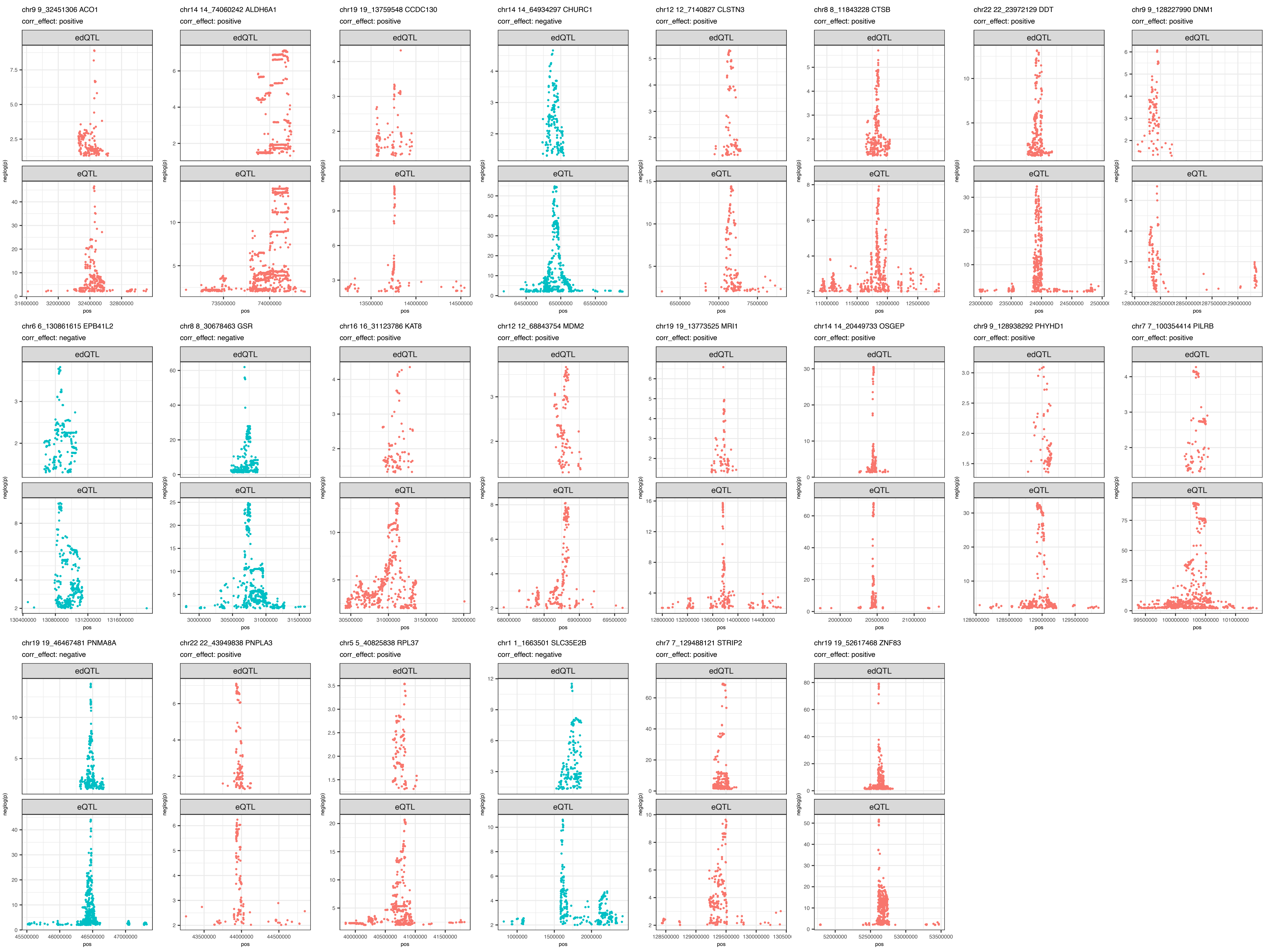
